# The PPE2 protein of *Mycobacterium tuberculosis* is responsible for the development of hyperglycemia and insulin resistance during tuberculosis

**DOI:** 10.1101/2025.08.13.670014

**Authors:** Manoj Kumar Bisht, Vandana Maurya, Priyanka Dahiya, Vijaya Lakshmi Valluri, Sudip Ghosh, Sangita Mukhopadhyay

**Author notes:** **To whom correspondence should be addressed:** Laboratory of Molecular Cell Biology, BRIC-Centre for DNA Fingerprinting and Diagnostics (CDFD), Inner Ring Road, Uppal, Hyderabad - 500039, Telangana, India.

## Abstract

Diabetes is a known risk factor for tuberculosis (TB), but clinical evidences suggest that TB itself can induce hyperglycaemia and insulin resistance, though the underlying mycobacterial factors are not known. Herein, we implicate PPE2, a secretory PE/PPE family protein of *Mycobacterium tuberculosis* (Mtb), as a key modulator of adipose tissue physiology that contributes to the development of insulin resistance. In mice, PPE2 caused fat loss, adipocyte hypertrophy, immune cell infiltration, impaired glucose tolerance, reduced expression of PPAR-γ, C/EBP-α, adiponectin and higher insulin resistance. Transcriptomic analysis revealed PPE2 altered expression of genes associated with chemokine/cytokine, ribosomal biogenesis and lipase signaling. PPE2 induced lipolysis by activating cAMP-PKA-HSL axis, increased circulating free fatty acids, a feature also observed in TB patient sera. Interestingly, PPE2-immunization mitigated these effects, suggesting its potential as a subunit vaccine. Overall, this study identifies PPE2 as a key link between Mtb-infection, adipose tissue dysfunction and insulin resistance.

## Introduction

*Mycobacterium tuberculosis* (Mtb) remains one of the deadliest bacterial pathogens, causing approximately 1.23 million deaths annually. Each year, over 10.7 million people are infected, but about 5-10% of them develop clinical symptoms of active tuberculosis (TB) disease (*Global Tuberculosis Report, 2025*). In the majority of cases, the bacilli enter a dormant phase during which individuals test positive on immunologic tests but do not exhibit clinical symptoms, a condition referred as latent tuberculosis (*Lin and Flynn, 2010*). Latent tuberculosis can reactivate into active tuberculosis when the immune system becomes compromised due to factors such as undernutrition, smoking, HIV co-infection, use of immunosuppressive drugs and diabetes (*Ai et al., 2016*). It has been found that people with diabetes are 1.5-2.4 times higher risk of developing TB (*Franco et al., 2024*) and diabetes is responsible for 10.6% of the total TB deaths (*GBD Tuberculosis Collaborators, 2018*). The co-occurrence of tuberculosis and diabetes (TB-DM) complicates TB treatment and drastically worsens disease outcomes. Several studies have reported that TB can exacerbate the development and severity of diabetes-associated complications (*Magee et al., 2018; Mugusi et al., 1990*). The TB-DM co-occurrence is a growing global health concern with an established bidirectional relationship between the two conditions (*Khattak et al., 2024; Gautam et al., 2021*). Diabetes is a well-known risk factor for tuberculosis with an estimated global prevalence of 15.3% among TB patients, varying by country of origin, age, sex, region, level of country income and development (*Noubiap et al., 2019*). The prevalence of diabetes in Nigeria (15%), Tanzania (11%), and Ethiopia (10%) (*Alebel et al., 2019*) is notable while in India it is as high as 25.3% (*Viswanathan et al., 2012*), which has the second largest number of diabetic patients in the world (*IDFD Atlas, 2025*).

Several clinical studies indicate that TB patients are prone to develop insulin resistance and hyperglycemia. In a Nigerian study, it has been found that about 37% of the patients with active pulmonary TB had impaired glucose tolerance, which could be reversed during anti-tuberculosis treatment (*Oluboyo and Erasmus, 1990*). More recently many studies have shown that TB patients with no known history of diabetes can develop hyperglycemia (*Magee et al., 2018; Lin et al., 2017*). Though transient hyperglycemia is commonly observed in many TB patients, 8-87% of TB patients are newly diagnosed with dysglycemia without a prior history (*Mave et al., 2017; Magee et al., 2018*).

Development of hyperglycemia, glucose intolerance and insulin resistance can generally be attributed to metabolic changes, induction of pro-inflammatory conditions in the host, and dysregulation of adipose tissue functions (*Guilherme et al., 2008*). Interestingly, Mtb is known to persist in the adipose tissue and to regulate pulmonary pathology and has been implicated in tuberculosis and changes in adipose tissue behavior (*Erol, 2008; Neyrolles et al., 2006; Beigier-bompadre et al., 2017*). Mtb persistence in adipose tissue is known to cause adipocyte hypertrophy, lipolysis, changes in physiology and infiltration of immune cells in adipose tissue (*Beigier-bompadre et al., 2017; Bisht et al., 2023a). Monocyte chemoattractant protein-1 (MCP-1) expression was found to be higher in mycobacterium-infected visceral adipose, which has been linked to increased immune cell infiltration, insulin resistance and hepatic steatosis (*Kanda et al., 2006*). Adipose tissue-resident Mtb is known to alter its physiology which may cause a shift in the metabolism of fatty acids (Neyrolles, 2006; Beigier-bompadre et al., 2017; Erol, 2008*).

Though direct interaction between Mtb and host adipose tissue exists, the nature and outcome of such interaction are not well understood. Therefore, we speculated that certain Mtb factors/proteins, which are either exposed on the cell wall or secreted into the host circulation or directly released into the adipose tissue milieu by resident Mtb, are likely to be involved in modulating adipose tissue physiology and functions. In this study, we investigated the role of a secretory mycobacterial protein, PPE2, in the modulation of adipose tissue physiology and functions. PPE2 belongs to the PE/PPE family and possesses a nuclear localization signal sequence and a leucine zipper DNA-binding motif, and is secreted during Mtb infection in mice (*Bhat et al., 2013; Bhat et al, 2017; Abraham et al., 2014; Bisht et al., 2023b*). PPE2 appears to be a pleiotropic protein and plays an important role in mycobacterial virulence (*Srivastava et al., 2019; Pal et al., 2021*). It inhibits nitric oxide (NO) production by binding to the GATA-1-like elements in the iNOS promoter, inhibits the production of reactive oxygen species (ROS) by destabilizing NADPH-oxidase complex, as well as inhibits myelopoiesis to reduce the production of myeloid cells like leukocytes, mast cells, platelets and red blood cells (RBCs) (*Bhat et al., 2017; Srivastava et al., 2019; Pal et al., 2021; Pal and Mukhopadhyay, 2021*).

To the best of our knowledge, for the first time, our study provides evidence that PPE2 can modulate gene expression and adipose tissue functions leading to glucose intolerance and hyperglycemia. Using recombinantly purified protein, a surrogate non-pathogenic mycobacterium expressing PPE2 and a *ppe2*-null mutant of a clinical *M. tuberculosis* strain (CDC1551), we have shown that PPE2 interferes with adipogenesis by inhibiting adipogenic transcription factors, inhibiting adipokines and promoting adipocyte hypertrophy and lipolysis, which can lead to the development of impaired glucose intolerance and insulin resistance.

## Results

### 1. Recombinant PPE2 protein (rPPE2) inhibits differentiation of adipocytes

*Mycobacterium tuberculosis* has been previously reported to inhibit adipogenic transcription factors like PPAR-γ and adipocyte secreted proteins like adiponectin, in the adipose tissue of infected mice (*Ayyappan et al., 2019*). PPE2 has been identified to be a secretory protein (*Bhat et al., 2013; Pal et al., 2022*), and its presence has been demonstrated in the bloodstream of mice infected with *M. smegmatis* and *M. tuberculosis* (*Bisht et al., 2023b*). Therefore, it is speculated that PPE2 may exert a systemic effect on the regulation of adipose tissue functions and thereby may contribute to diabetes-related complications. Therefore, to investigate the impact of PPE2 on adipogenesis, differentiation of 3T3-L1 pre-adipocyte was carried out in the presence of recombinantly purified PPE2 protein (rPPE2). Accordingly, 3T3-L1 cells were treated with a differentiation cocktail medium (*Sadowski et al., 1992*), in the absence or presence of various concentrations of rPPE2 and triglyceride accumulation was measured using Oil Red O staining at day 10 post-treatment. In the presence of rPPE2, the intensity of Oil Red O staining was found to be decreased in a concentration-dependent manner (Figure 1A, B), suggesting that PPE2 inhibits adipocyte differentiation. Significant inhibition was observed at rPPE2 concentration of 3 µg/ml, which was used for subsequent experiments. No cytotoxic effect of rPPE2 was observed when the 3T3-L1 preadipocytes were treated for 48 hours (Figure 1—figure supplement 1), suggesting that the observed inhibition of adipogenesis was due to physiological effect of rPPE2. To further confirm the specificity of inhibition, 3 µg/ml and 5 µg/ml of anti-PPE2 antibody (in house raised in Balb/c mice (*Bisht et al., 2023b*)) was used to neutralize the effect of rPPE2 during differentiation. It was found that, in the presence of anti-PPE2 Ab, the inhibitory effect of rPPE2 on adipogenesis was neutralised in a concentration-dependent manner (Figure 1C).

**Figure 1.**
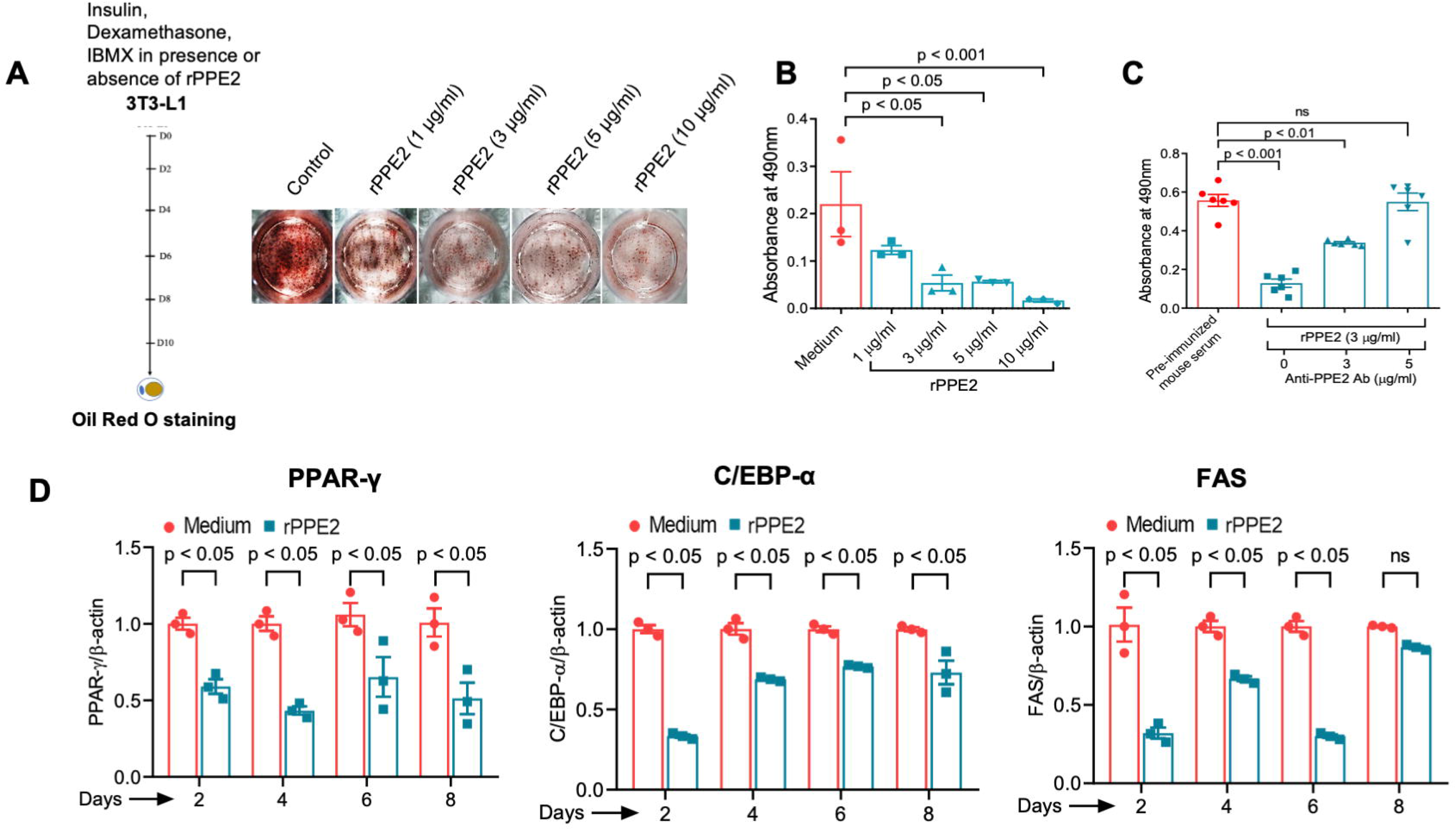
Recombinantly purified PPE2 protein of *M. tuberculosis* (rPPE2) inhibits differentiation of 3T3-L1 cells into adipocytes. **(A)** 3T3-L1 cells were treated with different concentrations of rPPE2 and induced to differentiate using DMI cocktail (dexamethasone, insulin and IBMX). At day 10 post-induction of differentiation, the accumulated triglycerides in adipocytes were stained with Oil Red O and image of differentiated adipocytes were taken. **(B)** For quantitative measurement of lipid accumulation, lipid-bound Oil Red O stain was extracted with isopropanol and absorbance of the extracted solution was measured at 490 nm using isopropanol as blank. **(C)** Differentiation of rPPE2-treated 3T3-L1 cells in the absence or presence of anti-PPE2 Ab. **(D)** 3T3-L1 cells were differentiated in the absence or presence of 3 µg/ml rPPE2 and changes in the expressions of PPAR-γ, C/EBP-α and FAS genes at various days post-differentiation were measured by qPCR. β-actin transcript level was used as an internal control. The results shown are mean ± SEM of 3 different experiments. ns = non-significant. Dose-dependent effects of rPPE2 and antibody-mediated rescue experiments were analyzed using one-way ANOVA followed by Dunnett’s test against control. Time-course experiments were analyzed using two-way ANOVA followed by Šídák test for each day. Statistical significance was considered when p < 0.05.

Next, the effect of PPE2 on adipocyte differentiation was examined in the context of mycobacterial infection. Accordingly, 3T3-L1 pre-adipocytes were infected with either *M. smegmatis* harbouring vector control (Msmeg-pVV) or *M. smegmatis* expressing PPE2 (Msmeg-PPE2), followed by induction of differentiation and Oil Red O staining. Cells infected with Msmeg-PPE2 exhibited a significant reduction in adipocyte differentiation compared to those infected with Msmeg-pVV (Figure 1—figure supplement 2), indicating that PPE2 could inhibit adipocyte differentiation when present in the context of bacilli. However, infection with Msmeg-pVV also caused a noticeable reduction in adipogenesis compared to uninfected control cells, indicating that other factors intrinsic to *M. smegmatis* or infection-triggered signaling may also contribute to the inhibition of adipogenesis. Next, we examined the effect of PPE2 on critical adipogenic factors like PPAR-γ and C/EBP-α and fatty acid synthase (FAS) during differentiation of 3T3-L1 cells. In the presence of rPPE2, there was a marked decrease in the expression of PPAR-γ, C/EBP-α and FAS during differentiation (Figure 1D). These findings suggest that PPE2 inhibits 3T3-L1 differentiation by downregulating key adipogenic genes involved in adipocyte differentiation, highlighting its potential role in modulating adipose tissue physiology and function.

### 2. PPE2 protein alters adipose tissue physiology during mice infection

Mtb can persist within adipose tissue and alter its physiology, potentially leading to a shift in the metabolism of fatty acids (*Neyrolles et al., 2006, Beigier-Bompadre et al., 2017, Erol, 2008*). While Mtb is known to cause inhibition of adipogenesis and promote lipolysis in adipose tissue (*Ayyappan et al., 2019*), the mechanisms responsible for this effect remained unclear. Since we found that PPE2 inhibits adipogenesis in 3T3-L1 cells, we investigated its impact on adipose tissue physiology and function during infection in mice. Balb/c mice of 9-11 weeks old were infected with either Msmeg-pVV or Msmeg-PPE2 and sacrificed on day 2 and day 7 post-infection. The two major fat depots, the perigonadal (PG) and the visceral (VS) fat tissues, were analysed to assess fat mass, tissue pathology, gene expression pattern and bacterial survival. Mice infected with Msmeg-PPE2 showed significant loss of adipose tissue mass (Figure 2—figure supplement 1) compared to both the uninfected control and Msmeg-pVV infected groups, indicating that PPE2 inhibits adipogenesis in both perigonadal and visceral fat depots (Figure 2A, B). Notably, loss of adipose tissue was also observed in Msmeg-pVV-infected mice, implying that either increased energy demands to combat infection or some intrinsic factors from *M. smegmatis* may also contribute to adipose tissue atrophy.

**Figure 2.**
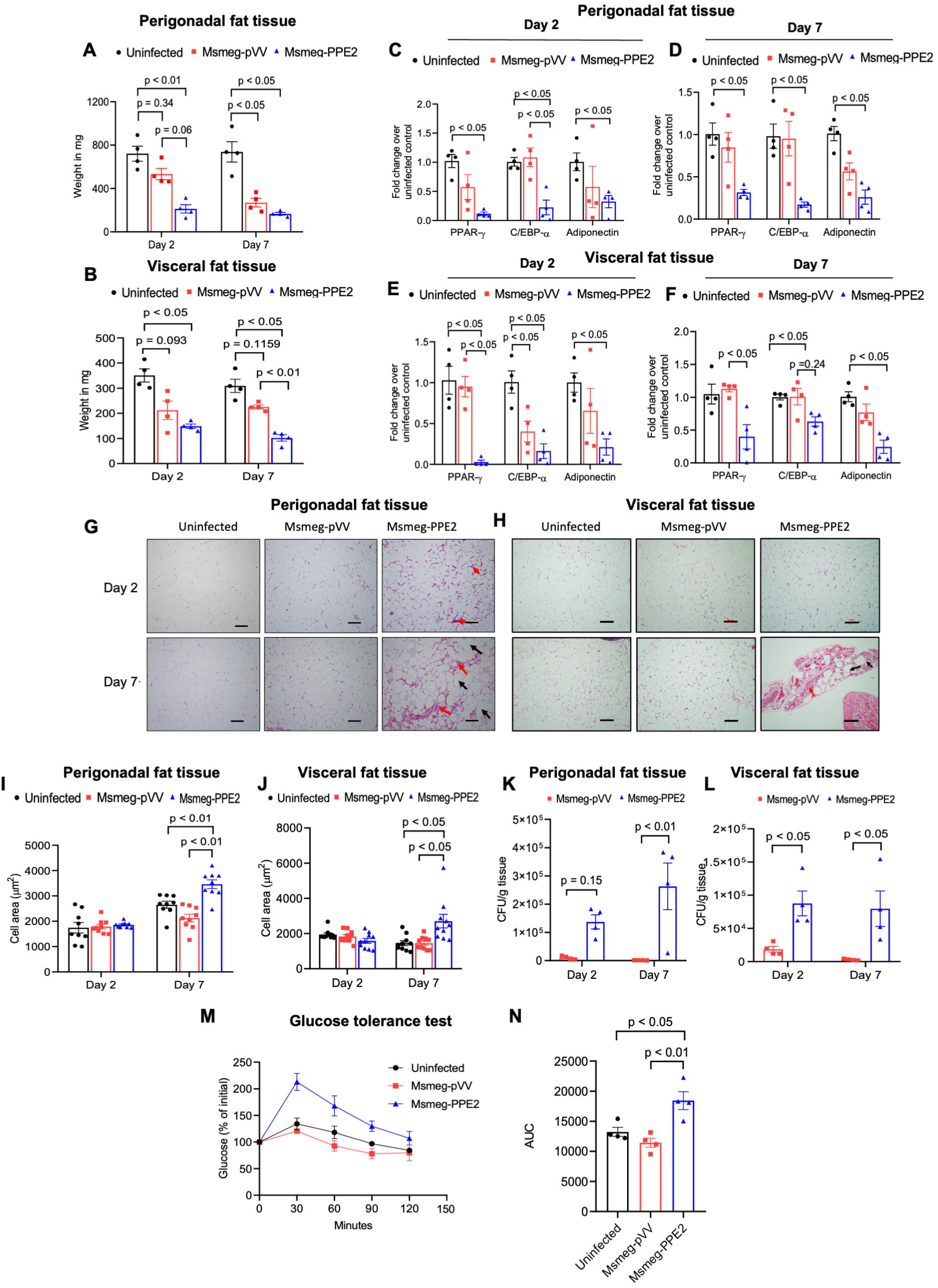
Infection of mice with *M. smegmatis* expressing PPE2 protein of *M. tuberculosis* alters adipose tissue physiology. **(A, B)** Balb/c mice were infected with 100 million of either Msmeg-pVV or Msmeg-PPE2 bacilli and sacrificed at 2^nd^ and 7^th^ day post-infection (dpi) to measure the weight of perigonadal (**A**) and visceral (**B**) fat tissue. **(C, D, E, F)** Expression levels of PPAR-γ, C/EBP-α and adiponectin in perigonadal tissue and in visceral tissue were quantified by qPCR at day 2 (**C** and **E**) and day 7 (**D** and **F**) post-infection (dpi). GAPDH transcript level was used as an internal control. **(G and H)** Histological analyses of perigonadal (**G**) and visceral (**H**) adipose tissue sections were performed at day 2 and day 7 post-infection by staining with hematoxylin and eosin (H&E) and photographs of representative sections are presented at 40X magnification (scale bar = 100 µm). Black arrows indicate adipocyte hypertrophy (enlarged adipocyte) and the red arrows point to the infiltrated immune cells into the adipose tissue. **(I and J)** Adipocyte cell area for perigonadal (**I**) and visceral (**J**) fat tissue was measured using ImageJ software. **(K and L)** Bacterial load in perigonadal (**K**) and visceral (**L**) fat tissue was measured at day 2 and day 7 post-infection. For **(M and N)** For glucose tolerance test at day 7 post-infection, mice were fasted overnight followed by intraperitoneal injection of glucose (2 g/kg body weight) and blood glucose levels were monitored for 120 minutes (**M**) and the Area under the curve (AUC) was calculated (**N**). The results shown are mean ± SEM of 4 mice. For figures A-F and I-J, two-way ANOVA followed by Tuckey’s HSD test were carried out. For K-L, two-way ANOVA followed by Tukey’s test was carried out. For figure N, one-way ANOVA followed by Tukey’s HSD test was carried out. p value of < 0.05 was considered to be significant.

Next, we examined the expression patterns of PPAR-γ, C/EBP-α, as well as adiponectin in these infected mice. PPAR-γ is a key transcription factor involved in regulating adipocyte differentiation, lipid metabolism and insulin sensitivity (*Maréchal et al., 2018*), while C/EBP-α is also required for adipogenesis as well as normal adipocyte function, while adiponectin is known to play important roles in variety of physiological functions including lipid metabolism, energy homeostasis, inflammation, and insulin sensitivity (*Khoramipour et al., 2021*). Infection with Msmeg-PPE2 resulted in a marked decrease in the expression of PPAR-γ, C/EBP-α and adiponectin in both PG and VS adipose tissues compared to mice infected with Msmeg-pVV at both 2^nd^ and 7^th^ day post-infection (dpi) (Figure 2C-F). These findings suggest that PPE2 during mycobacterial infection leads to altered gene expression, which potentially contributes to the modulation of the physiology and function of adipose tissue.

Adipose tissue is composed of various cell types including preadipocytes, adipocytes, endothelial cells, blood cells, fibroblasts, macrophages and several types of other immune cells (*Saetang and Sangkhathat, 2018; Richard et al., 2020*). During mycobacterial infection, immune cells infiltrate into adipose tissue and play a role in regulation of inflammation (*Beigier-Bompadre et al., 2017*). Histological analyses of adipose tissues stained with H&E revealed that mice infected with Msmeg-PPE2 exhibited increased immune cell infiltration and adipocyte hypertrophy by 7^th^ dpi (Figure 2G and H). Higher infiltration of immune cells as well as adipose tissue hypertrophy was found to be more pronounced at 7^th^ dpi in both the fat depots. Notably, at 7^th^ dpi, an increased cell surface area was observed in perigonadal and visceral fat tissues of Msmeg-PPE2-infected mice in comparison to uninfected or Msmeg-pVV-infected mice (Figure 2I and J). These findings collectively indicate that Msmeg-PPE2 infection leads to enhanced immune cell infiltration and adipose tissue hypertrophy, influencing the cellular composition and morphology of adipocytes. We also assessed mycobacterial survival within perigonadal and visceral fat tissues. Adipose tissues from mice infected with Msmeg-PPE2 had a higher colony forming unit (CFU) count as compared to those infected with Msmeg-pVV at both 2^nd^ and 7^th^ dpi (Figure 2K and L). This suggests that PPE2 contributes to enhanced survival of mycobacteria within PG and VS adipose tissues. These data suggest that PPE2 can significantly influence the adipose tissue at the morphological and molecular levels. Since adipose tissue plays a crucial role in regulating immune metabolism and glucose homeostasis, an intraperitoneal glucose tolerance test (IPGTT) was performed in mice at 7^th^ dpi. After overnight fast, all mice received an intraperitoneal injection of dextrose (2 g/kg body mass), and blood glucose levels were monitored over 120 minutes. Interestingly, mice infected with Msmeg-PPE2 exhibited increased glucose intolerance in comparison to both uninfected and Msmeg-pVV-infected mice at 7^th^ dpi (Figure 2M). Analyses of the area under the curve (AUC) indicate that Msmeg-PPE2-infected mice had higher glucose intolerance compared to other groups (Figure 2N). These findings indicate that PPE2 may impair glucose homeostasis through its effects on adipose tissue physiology.

### 3. Administration of recombinantly purified PPE2 protein (rPPE2) causes metabolic perturbations in mice

In the previous section, it was found that PPE2, when presented in the context of bacterium can modulate adipose tissue physiology in mice. To further confirm that the observed changes can be directly attributed to the PPE2 protein, mice were injected intraperitoneally with various concentrations of rPPE2 (100 μg/kg, 300 μg/kg, 1 mg/kg, 3 mg/kg) or bovine serum albumin (BSA) as a control (1 mg/kg) on alternate days for 12 days. On morphological examination of PG and VS fat depots, a dose dependent reduction in both size and mass was observed in rPPE2-administered mice compared to those administered with BSA and 3 mg/kg showed the highest decrease (Figure 3A-C). To assess the impact of rPPE2 on glucose homeostasis, an IPGT test was performed and the results indicate a significant metabolic effect, with 3 mg/kg rPPE2-treated mice (but not in the treatment groups received 100 μg/kg, 300 μg/kg and 1 mg/kg rPPE2 protein) displaying a significantly lower rate of glucose clearance as compared to the control mice treated with BSA (Figure 3D and E). This observation indicates that PPE2 is a potential mycobacterial factor in the development of hyperglycaemia during tuberculosis.

**Figure 3.**
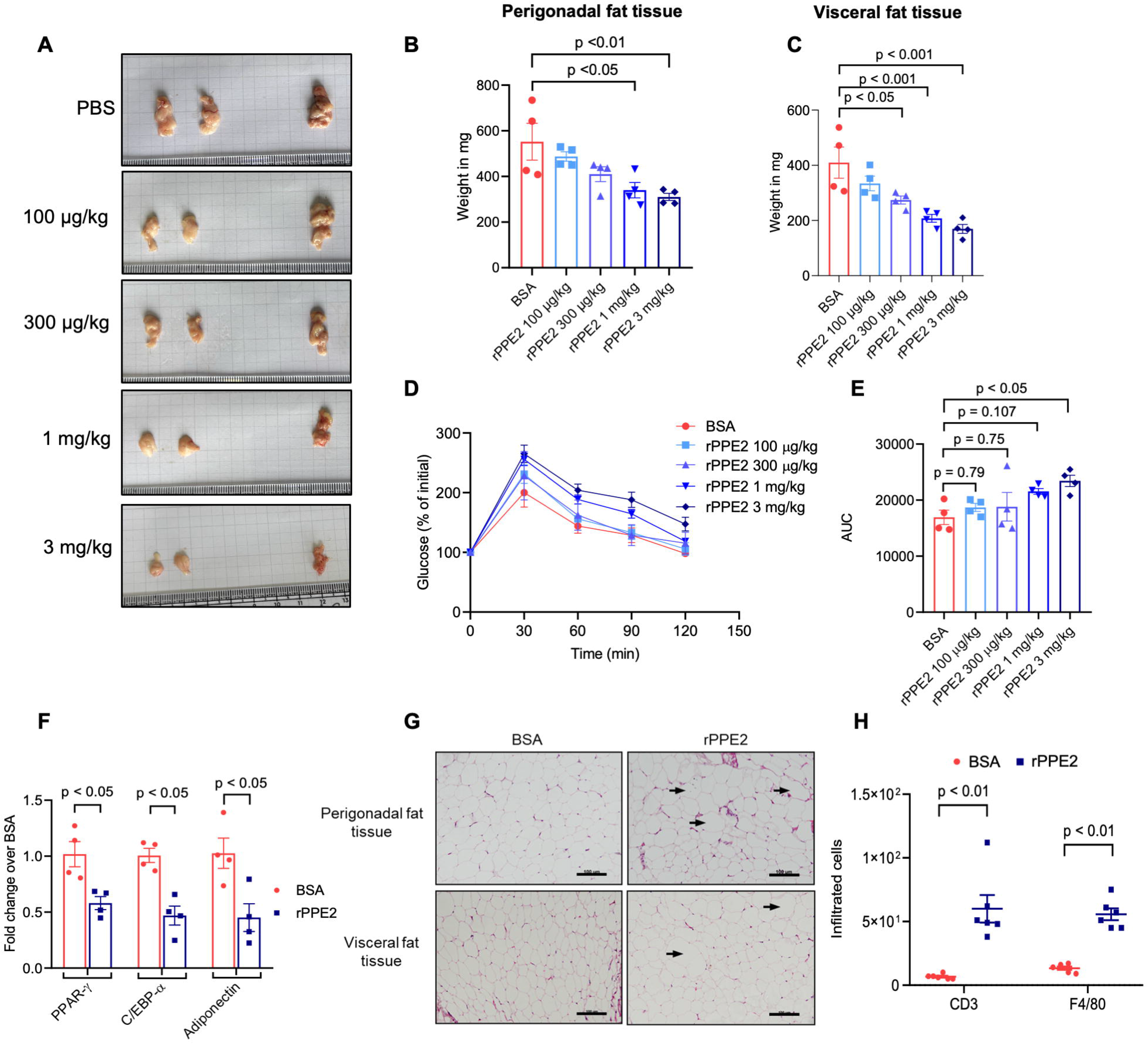
Administration of rPPE2 protein leads to adipose tissue loss and impaired glucose tolerance in mice. Balb/c mice were intraperitoneally administered with 3 mg/kg of either BSA or recombinant PPE2 protein (rPPE2) on alternate days for 12 days and mice were sacrificed at day 14 post-treatment. **(A)** Representative photographic images of perigonadal and visceral fat tissues are shown. (**B and C**) Weight of the total perigonadal fat mass (**B**) and visceral fat mass (**C**) are recorded. (**D and E**) Mice were starved overnight and glucose tolerance test was performed for 120 minutes at day 14 post-treatment **(G)** and the mean Area under the curve (AUC) was calculated (**E**). (**F**) Transcript levels of PPAR-γ, C/EBP-α and adiponectin genes are quantified in perigonadal fat tissue using qPCR where β-actin transcript level was used as internal control. **(G)** Histopathology analyses of perigonadal and visceral fat tissue (representative photographs of hematoxylin and eosin (H&E)-stained sections) is shown at 40X magnification (scale bar = 100 µm). Black arrows indicate adipocyte hypertrophy (enlarged adipocyte) and the red arrows indicate infiltrated immune cells into the adipose tissue. (**H**) Number of infiltrated CD3^+^ T-cells and F4/80^+^ macrophages were counted using ImageJ software and plotted as cells per field of view for PG. The results shown are mean ± SEM of 4 mice. For figures B-E, one-way ANOVA followed by Dunnet’s test was carried out. For figures F and H, Student’s *t* test was carried for each gene/marker. p value of < 0.05 was considered to be significant.

Next, the histopathology and gene expression pattern were analysed for the group received 3 mg/kg rPPE2 to assess the direct impact of PPE2 protein on adipose tissue physiology. Importantly, 3 mg/kg rPPE2-administration led to significant decrease in the expression of PPAR-γ, C/EBP-α and adiponectin in the PG fat tissue in comparison to those treated with BSA (Figure 3F). These data indicate that PPE2 impair expression of key genes involved in adipogenesis and immune-metabolic functions. Histopathological studies of PG and VS fat tissues in rPPE2-treated mice revealed pronounced adipocyte hypertrophy and increased infiltration of immune cells into the adipose tissue marked by intense staining of the nucleus of the infiltrated cells (Figure 3G). Since increased infiltration of immune cells like T-cells, macrophages (*Wu et al., 2007*) into the adipose tissue have been implicated in adipocyte dysfunction and development of insulin resistance (*Apovian et al., 2008; Guilherme et al., 2008*), we next compared the degree of these immune cell infiltration in PG and VS adipose tissue of mice treated with 3 mg/kg rPPE2 *versus* BSA. The analysis of immune cell infiltration in these adipose tissues using anti-CD3^+^ antibody (Ab) for staining T-cells and anti-F4/80⁺ Ab for staining macrophages, revealed increased infiltration of both T-cells and macrophages in the rPPE2-treated group compared to BSA-treated group, indicative of adipose tissue inflammation and metabolic dysfunction by PPE2 protein (Figure 3H).

Since PPE2 is a secretory protein and detectable in the circulation of Mycobacterium-infected mice (*Bisht et al., 2023b*) and capable of promoting glucose intolerance, we next sought to determine whether neutralization of circulating PPE2 could ameliorate PPE2-induced glucose intolerance during mycobacterial infection. Accordingly, mice were immunized with rPPE2 using Freund’s incomplete adjuvant, followed by three booster doses at 15-day intervals. All the immunized mice showed a higher titre of anti-PPE2 antibody in the sera (Figure 2—figure supplement 2). Subsequently, these mice were infected with either Msmeg-pVV or Msmeg-PPE2, while mice injected with PBS served as unimmunized control group. As expected, loss of PG and VS fat tissues was effectively prevented in Msmeg-PPE2 infected mice which were previously immunized with rPPE2 on 7^th^ dpi (Figure 2—figure supplement 3A-C). This is reflected in significant improvement in the total body fat of the Msmeg-PPE2-infected mice which were immunized previously. In contrast, unimmunized mice showed significant loss of total body fat content (Figure 2—figure supplement 3D). Histopathological examination of both PG and VS adipose tissues showed marked improvement in the pathology in mice immunized with rPPE2 when compared to unimmunized mice (Figure 2—figure supplement 3E). Moreover, reduced infiltration of cells was observed in PPE2-immunized mice compared to unimmunized mice infected with Msmeg-PPE2 (Figure 2—figure supplement 3E). The rPPE2-immunized mice also exhibited significantly improved glucose tolerance as revealed by IPGT test (Figure 2—figure supplement 3F and G). In summary, rPPE2-immunized mice showed marked improvement in adipose tissue content, total body fat content, tissue pathology and glucose tolerance when challenged with infection, underscoring the role of PPE2 in modulating adipose tissue physiology and function.

### 4. PPE2 alters adipose tissue physiology and induces glucose intolerance in mice during infection with *M. tuberculosis*

The role of adipose tissue in regulating pulmonary pathology, inflammation and *M. tuberculosis* load in murine TB model is now well established (*Ayyappan et al., 2019*). Disruptions in the metabolic equilibrium within the adipose tissue and/or dysfunction of adipocytes during Mtb infection can contribute to the development of insulin resistance (*Oswal et al., 2022*). In several clinical studies, stress-induced hyperglycemia has been reported in tuberculosis patients which tend to normalize with proper management (*Magee et al., 2018*). Moreover, increased prevalence of impaired glucose tolerance and diabetes appears to be common among the TB patients (*Oluboyo and Erasmus, 1990; Jeon et al., 2010; Mugusi et al., 1990; Raghuraman et al., 2014*). However, the exact etiological mechanisms responsible for the development of glucose intolerance and insulin resistance in TB patients are not yet clearly understood.

In the previous sections, we demonstrated that the PPE2 protein of *M. tuberculosis* can significantly remodel the adipose tissue and can cause significant impaired glucose tolerance in mice. To further confirm its pathophysiological effects, we used a *ppe2-*null mutant of a clinical Mtb strain CDC1551 (CDC1551-ΔPPE2) and compared its effect with the wild-type strain (CDC1551) and a *ppe2*-complemented strain (CDC1551-ΔPPE2::PPE2). Accordingly, 9-10 weeks Balb/c mice were aerosol-infected either with CDC1551 (wild-type) or *ppe2*-null mutant CDC1551 or CDC1551-ΔPPE2::PPE2 and the infected mice were sacrificed after 30-dpi and 60-dpi. Expectedly, a significant loss of adipose tissue mass was observed in both PG (Figure 4A) and VS (Figure 4B) fat depot in mice infected with the wild-type CDC1551 and the *ppe2*-complemented strains as compared to uninfected controls and mice infected with *ppe2*-null mutant (Figure 4A, B Figure 4—figure supplement 1), suggesting impairment of normal adipogenesis. Histological analyses revealed enhanced infiltration of immune cells into both PG and VS adipose tissues of the mice infected with wild-type Mtb as well as *ppe2*-complemented strain as compared to uninfected or CDC1551-ΔPPE2 strain-infected mice at both time points (Figure 4C). While adipocyte hypertrophy was notably conspicuous in the fat tissues of both the wild-type and *ppe2*-complmented strain-infected mice, CDC1551-ΔPPE2-infected mice also exhibited some degree of hypertrophy in adipocytes as compared to those of uninfected controls (Figure 4D, E).

**Figure 4.**
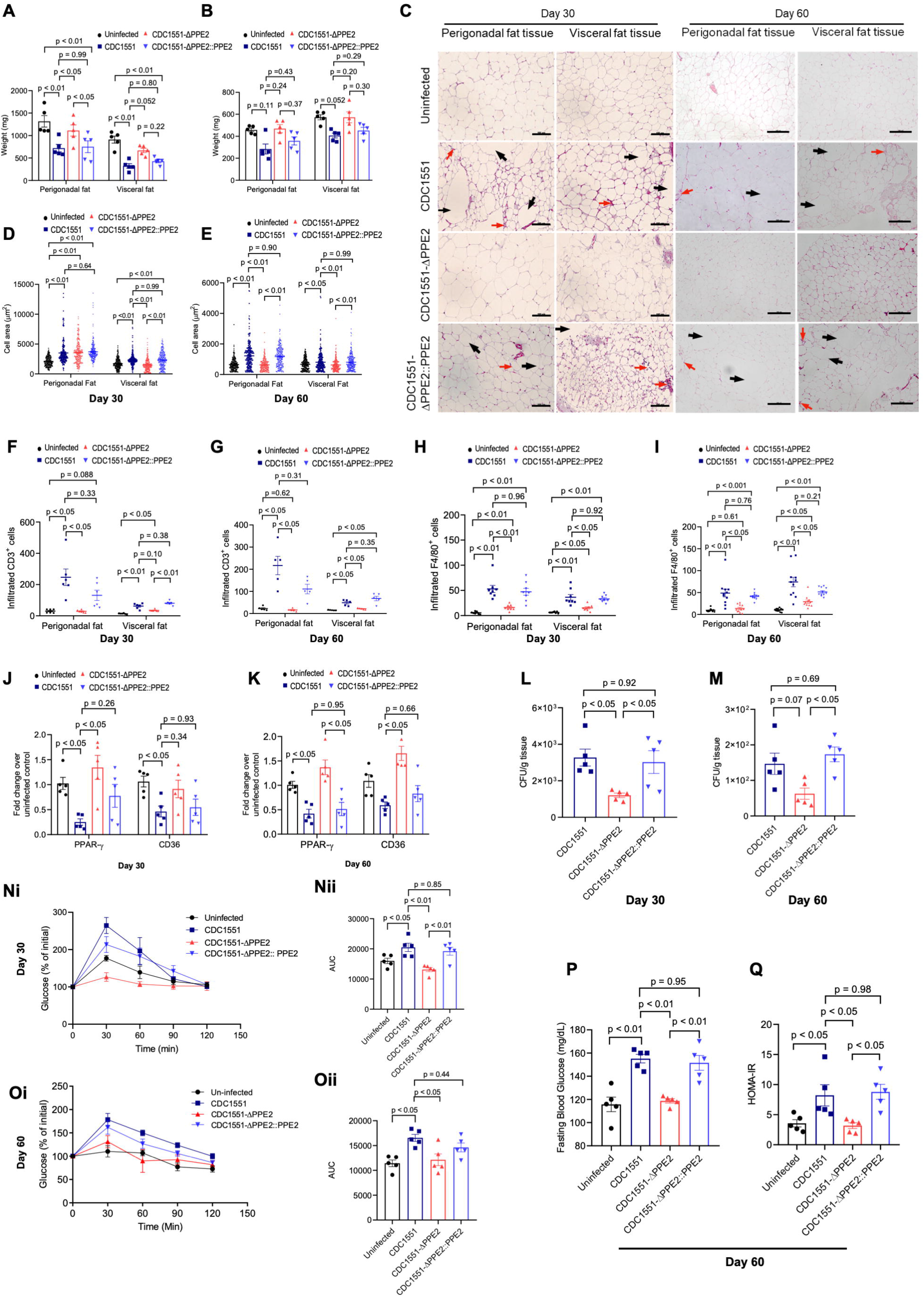
PPE2 alters adipose tissue physiology during *M. tuberculosis* infection in mice. Balb/c mice were infected through aerosol route either with *M. tuberculosis* clinical strain CDC1551 (wild-type) or CDC1551-ΔPPE2 or CDC1551-ΔPPE2::PPE2 (complemented) strains and were sacrificed at day 30 (**A**) and day 60 (**B**) post-infection (dpi). Perigonadal (PG) and visceral (VS) fat tissues were harvested and weight of PG fat tissue (**A**) and VS fat tissue (**B**) was measured. (**C**) Also, representative photographs of hematoxylin and eosin (H&E)-stained sections of PG at both day 30 and day 60 post-infection were visualized at 40X magnification (scale bar = 100 µM). Black arrow shows adipocyte hypertrophy (enlarged adipocyte) and red arrow represent infiltrated immune cells into the adipose tissue. (**D-E**) Graphs showing cell area of adipocytes in PG and VS fat tissues at 30 dpi (**D**) and 60 dpi (**E**) respectively. (**F-I**) Number of infiltrated CD3^+^ T-cells and F4/80^+^ macrophages were counted using ImageJ software and plotted as cells per field of view for PG and VS adipose tissue sampled at 30 dpi (**F and H**) and 60 dpi (**G and I**) respectively. (**J and K**) Changes of expression of PPAR-γ and CD36 in perigonadal tissue at day 30 (**J**) and day 60 (**K**) post-infection were measured by qPCR and β-actin transcript level was used as an internal control. (**L and M**) Number of bacilli in PG fat tissue at 30 (**L**) and 60 (**M**) dpi respectively. (**N and O**) IPGTT performed on mice along with corresponding AUC data obtained at day 30 (**Ni, Nii**) and day 60 (**Oi**, **Oii**) post-infection, respectively. (**P**) Fasting blood glucose levels of mice at 60 dpi and (**Q**) HOMA-IR scores in each mice group were calculated at day 60 post-infection. The results shown are mean ± SEM of 5 mice. For figures A-B and C-K, two-way ANOVA followed by Tuckey’s HSD test was carried out. For figures L-Q, one-way ANOVA followed by Tukey’s HSD was carried out. p value of < 0.05 was considered to be significant.

Mtb is known to promote infiltration of T-cells, macrophages cells causing impaired adipose tissue functions (*Beigier-Bompadre et al., 2017*). Therefore, we next examined the degree of immune cell infiltration in the PG and VS adipose tissues in mice infected with various mycobacterial strains at day 30 and day 60 post-infection using immunofluorescence microscopy. Significantly increased infiltrations of CD3^+^ T-cell and F4/80^+^ macrophage was observed in PG and VS adipose tissues of the mice infected with wild-type and *ppe2-*complemented Mtb strains as compared to those observed in *ppe2-*null mutant strain-infected mice at both 30 and 60 dpi (Figure 4F-I; Figure 4—figure supplement 2).

We also examined the changes in the expression of key genes associated with the maintenance and regulation of adipose tissue, such as PPAR-γ and CD36 (a scavenger receptor that is regulated by PPAR-γ and is involved in the uptake of long-chain fatty acids and other lipids) (*Maréchal et al., 2018; Chen et al., 2022*) in the perigonadal fat tissue of these infected mice. It was found that expression of both PPAR-γ and CD36 was significantly downregulated in mice infected with wild-type and *ppe2*-complemented strains, in comparison to mice infected with CDC1551-ΔPPE2 or uninfected control (Figure 4J, K). These results reiterate our earlier observations and underscore that PPE2 interferes with adipogenesis by suppressing the expression of adipogenic genes like PPAR-γ. Since, adipose tissue was previously speculated to be as an important dormancy niche for Mtb (*Neyrolles et al., 2006; Ayyappan et al., 2019*), we assessed PPE2’s impact on the survival of Mtb in adipose tissue particularly in perigonadal adipose tissue, for being a major reservoir of fat in mice. It was found that perigonadal adipose tissue from mice infected with wild-type as well as *ppe2*-complementated strains exhibited significantly higher bacterial load compared to CDC1551-ΔPPE2 strain-infected mice (Figure 4L, M). These data suggest that PPE2 promotes persistence of Mtb in adipose tissue.

We also assessed the impact of PPE2 on the rate of glucose clearance using IPGTT assay and degree of insulin resistance using homeostatic model assessment of insulin resistance (HOMA-IR) (*Matthews et al. 1985*), where the conversion factor was adapted from *Knopp et al. (2019)*. As expected, mice infected with the wild-type and *ppe2*-complemented strains exhibited markedly higher glucose intolerance compared to uninfected control mice and those infected with the CDC1551-ΔPPE2 strain at both 30 and 60 dpi, as indicated by blood glucose levels and AUC at 30-minute intervals (Figure 4Ni, 4Nii, 4Oi, 4Oii). At day 60 post infection, groups infected with wild-type Mtb and CDC1551-ΔPPE2 strain complemented with a functional *ppe2* gene showed higher fasting blood glucose and HOMA-IR compared to the groups uninfected or infected with CDC1551-ΔPPE2 strain (Figure 4P, Q). Therefore, these data indicate that mycobacterial infection can influence glucose homeostasis leading to the development of early hyperglycaemia which can later lead to development of insulin resistance and PPE2 is one of the potential etiological factors responsible for such metabolic perturbations.

### 6. PPE2 modulates the transcriptome of perigonadal adipose tissue

Results obtained from the experiments carried out in the earlier sections indicate that PPE2 can inhibit adipogenesis by modulating expressing of key adipogenic genes required for adipogenesis. To further assess its impact on the global gene expression profile in the adipose tissue, we performed RNA-seq in perigonadal adipose tissue collected from uninfected, wild-type and CDC1551-ΔPPE2-infected mice at 60 dpi. Compared to the uninfected, 3569 and 2441 genes were found to be differentially expressed in wild-type and CDC1551-ΔPPE2-infected mice adipose tissue, respectively, while 1261 genes were differentially expressed in CDC1551-ΔPPE2 and wild-type adipose tissue (Figure 5A). However, with ± 1.3 log2 fold change cut-off, we observed 807 and 661 genes differentially expressed in wild-type and CDC1551-ΔPPE2 adipose tissue, respectively, and 191 genes between CDC1551-ΔPPE2 and wild-type adipose tissues (Figure 5B, C). In significantly modulated DEGs among CDC1551-ΔPPE2 and wild-type infected mice, 96 genes overlapped with mice infected with wild-type Mtb, and 76 were unique DEGs in CDC1551-ΔPPE2- and wild-type-infected mice adipose tissues. To identify key signaling pathways influenced by PPE2 in adipose tissue, we performed gene set enrichment analysis (GSEA) using gene ontology, KEGG, and REACTOME pathway databases (*Subramanian et al., 2005*) using the DEGs between wild-type- and CDC1551-ΔPPE2-infected mice (Figure 5D and E). As expected, gene sets associated with tissue-specific immune responses (chemokine and cytokine signaling) were highly upregulated in mice infected with wild-type Mtb (Figure 5F and Figure 5—figure supplement 1), while gene sets related to oxidative phosphorylation were found to be downregulated in these mice (Figure 5F). Phospholipase C activity was found to be highly enriched in the adipose tissue of wild-type CDC1551 compared to uninfected and CDC1551-ΔPPE2-infected mice, as represented by the genes *TXK, CD86, GPR55*, *Angiotensin*, and *LPAR1* (Figure 5G). Both gene ontology and Reactome pathway analyses showed enrichment of genes related to cytosolic ribosomal protein synthesis and ribosomal biogenesis in the adipose tissue of wild-type Mtb-infected mice compared to that of CDC1551-ΔPPE2-infected mice (Figure 5D, E and Figure 5H). Genes related to cytoplasmic translation of both small and large ribosomal subunits were found to be enriched in the adipose tissue of CDC155-infected mice. In addition, KEGG pathways analysis revealed enrichment of pathways related to prion disease, Alzheimer, Parkinson’s disease and other neurodegenerative pathways suggesting an increase in the unfolded proteins in the endoplasmic reticulum (ER). ER stress is known to trigger inflammation and lipolysis in adipocytes leading to development of dyslipidemia and insulin resistance (*Khan and Wang, 2014; Foley et al., 2021; Deng et al., 2012*). To validate the occurrence of ER stress, we performed real-time PCR analysis to assess the expression of ER stress markers like ATF4, a transcription factor that regulates stress response and GRP78, a chaperone which facilitates protein folding during unfolded protein response (UPR). It was found that the expression of both ATF4 and GRP78 was significantly upregulated in the perigonadal adipose tissue of wild-type Mtb-infected mice as compared to uninfected and CDC1551-ΔPPE2 mice groups, confirming induction of ER stress in the adipose tissue (Figure 5I and J).

**Figure 5.**
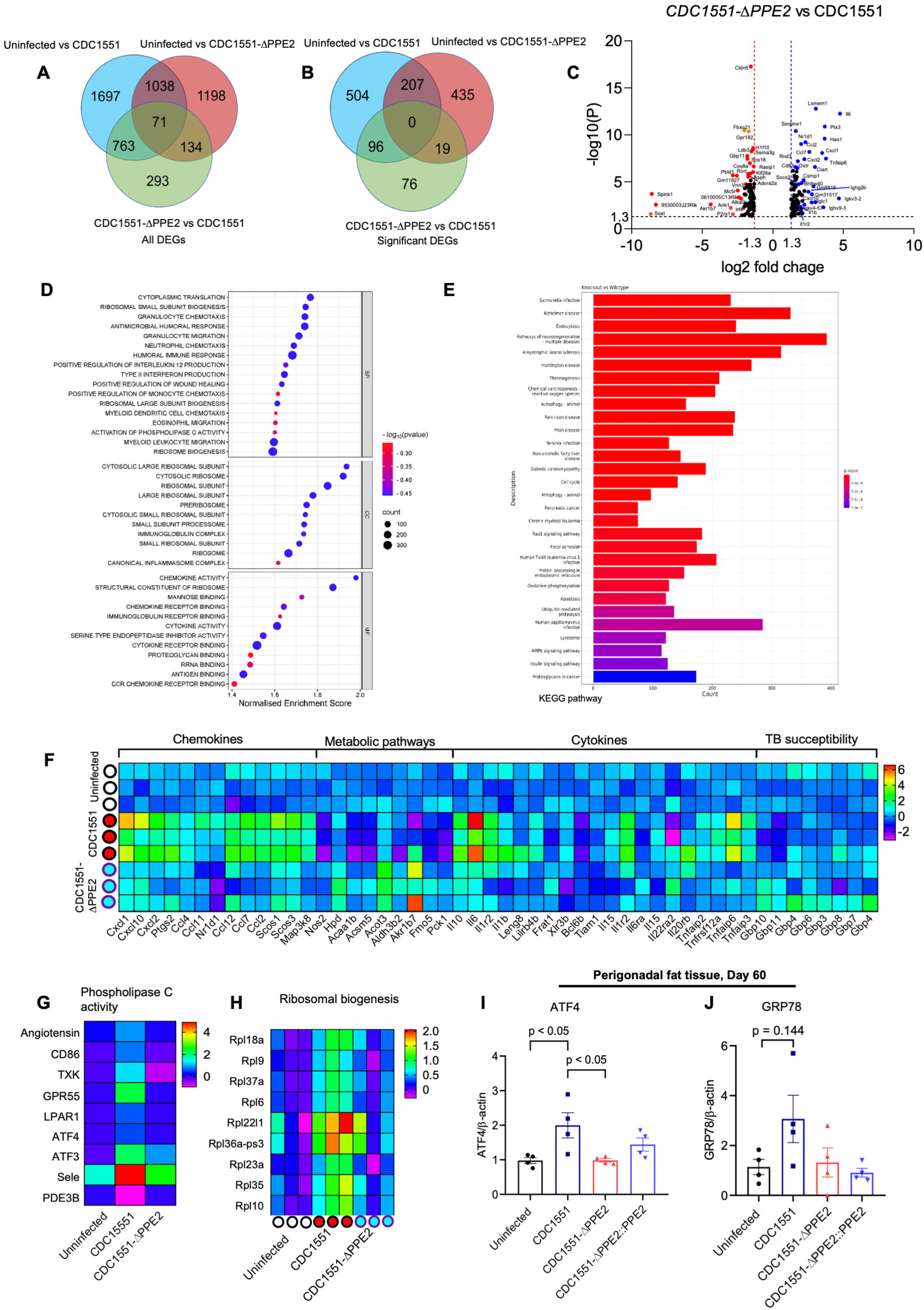
PPE2 modulates the adipose tissue transcriptome during infection. Perigonadal fat tissues were harvested from mice (n = 3) infected with either wild-type or CDC1551-ΔPPE2 strains or left uninfected at 60 dpi and subjected to RNA sequencing using Illumina platform. (**A**) Venn diagram showing all the differentially expressed genes (DEGs) in all subsets (**B**) Venn diagram showing significant DEGs after applying ±1.3-fold change cutoff. (**C**) Volcano plot showing significant DEGs in CDC1551-ΔPPE2 *versus* wild-type-infected mice in PG adipose tissue. (**D**) Gene set enrichment analysis (GSEA) for gene ontology analysis (**E**) KEGG pathway analysis in CDC1551-ΔPPE2 *versus* CDC1551 adipose tissues. (**F**) Heat map of DEGs for chemokine, metabolic pathways, cytokines and TB susceptibility genes in PG adipose tissues. (**G**) Heat map showing differential expression of phospholipase C activity genes. (**H**) Heat map showing differentially expressed ribosomal biogenesis genes. (**I and J**) Perigonadal fat tissue were harvested from mice either left uninfected or infected with various strains of *M. tuberculosis* at day 60 post-infection and used for cDNA synthesis. Transcript levels of ATF4 (**I**) and GRP78 (**J**) as measured by qPCR with β-actin as an internal control. The results shown are mean ± SEM of 4 mice. For figures I-J, one-way ANOVA followed by Tukey’s HSD test was carried out. p value of < 0.05 was considered to be significant.

### 8. PPE2 induce lipolysis during Mtb infection in adipocyte cell lines and in *vivo*

Since, ER stress in adipose tissue is known to trigger lipolysis through activation of cAMP/PKA pathway (*Song et al., 2017; Deng et al., 2012*), we next examined whether PPE2 can trigger lipolysis in mature 3T3-L1 adipocytes *in vitro* by measuring free glycerol released in the medium due to hydrolysis of triglycerides (*Schweiger et al., 2014*). Treatment of mature 3T3-L1 adipocytes with rPPE2 for 24 hours resulted in a dose-dependent increase of free glycerol in the medium (Figure 6A). Infiltration of immune cells and hypertrophy are known to stimulate production of pro-inflammatory cytokines such as TNF-α, which in turn is also known to trigger lipolysis (*Samad et al., 1999*). To determine whether PPE2-induced lipolysis occurs independently of TNF-α production, we treated mature 3T3-L1 adipocytes with a TNF-α inhibitor in the presence of rPPE2. While the inhibitor was able to effectively suppress TNF-α-induced free glycerol release, it did not significantly affect lipolysis induced by rPPE2 (Figure 6B). Since insulin possess antilipolytic properties (*Arner et al., 1981*), we next examined whether PPE2 can override antilipolytic effect of insulin and it was found that insulin failed to suppress rPPE2-induced lipolysis in 3T3-L1 adipocytes (Figure 6—figure supplement 1). Therefore, we speculated that PPE2 interferes with insulin’s ability to activate phosphodiesterase activity in adipocytes. To confirm this, we analysed the expression of phosphodiesterase 3B (PDE3B), a critical enzyme that converts cyclic adenosine monophosphate (cAMP) into 5’AMP and serves as a primary mediator of insulin’s antilipolytic effects (*Kitamura et al., 1999; DiPilato et al., 2015*). Semi-quantitative PCR analysis revealed that PPE2 treatment could significantly downregulate PDE3B expression in mature adipocytes (Figure 6Ci, 6Cii). Reduction in PDE3B is associated with elevated cAMP levels, which activate cAMP-dependent protein kinase (PKA) and activated PKA phosphorylates hormone-sensitive lipase (HSL), a key enzyme involved in lipolysis (*Degerman et al., 1998*). To confirm the involvement of this pathway, we assessed HSL phosphorylation levels in rPPE2-treated adipocytes at 6- and 24-hour post-treatment. The results showed a significant increase in HSL phosphorylation, indicating enhanced enzymatic activity (Figure 6Di, 6Dii). Furthermore, pretreatment with H89-dihydrochloride, a PKA-specific inhibitor, significantly reduced free glycerol release in rPPE2-treated adipocytes (Figure 6E). Collectively, these data suggest that PPE2 promotes lipolysis in adipocytes by reducing PDE3B expression, and activating PKA-HSL pathway through elevated cAMP levels.

**Figure 6.**
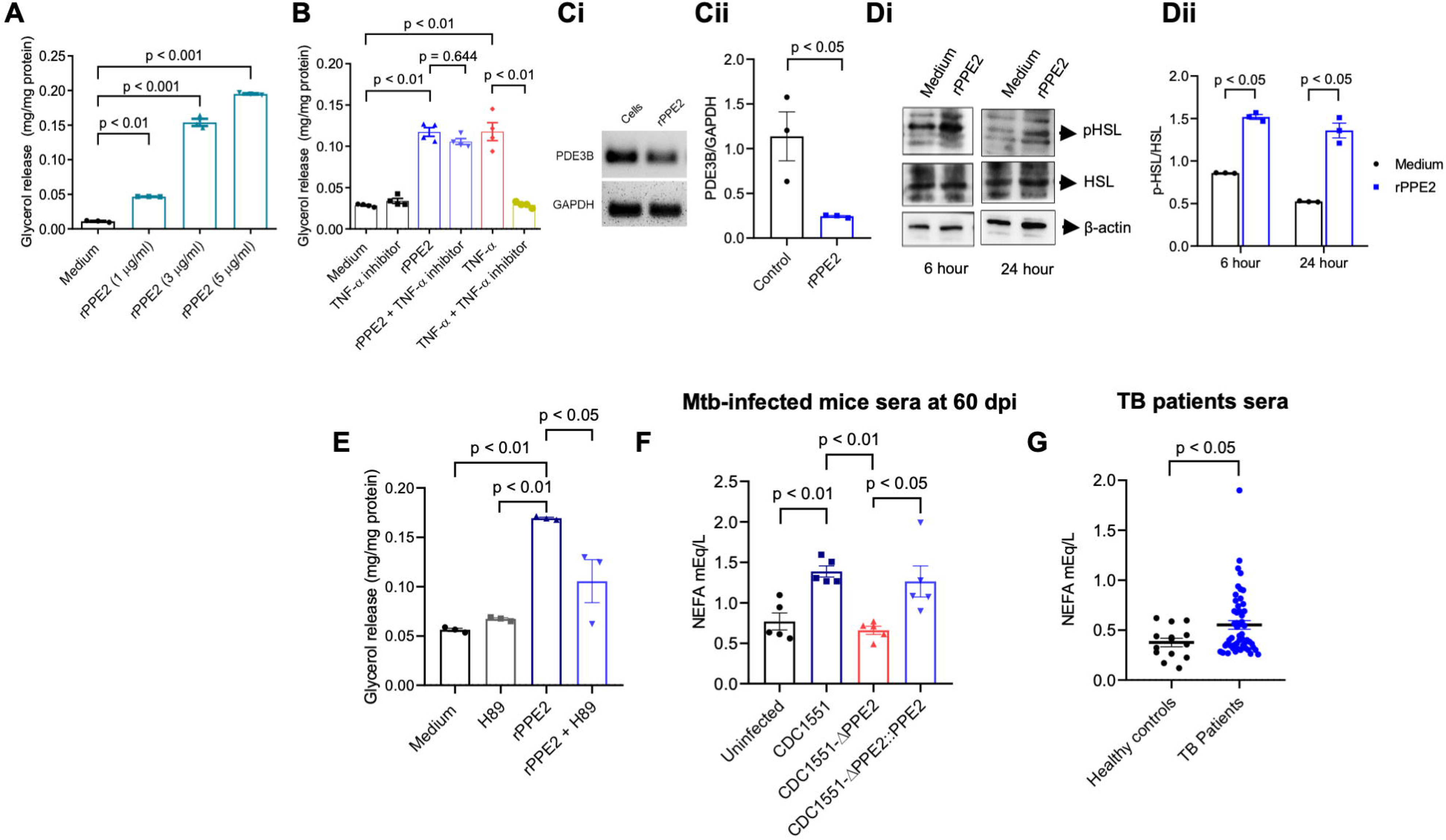
PPE2 induces lipolysis in adipocytes by targeting the cAMP-PKA-HSL signaling axis. (**A**) 3T3-L1 cells were differentiated into mature adipocytes and then treated with various concentration of rPPE2 and after 24 hours medium was removed from cells and amount of glycerol released was quantified. (**B**) 3T3-L1 mature adipocytes were incubated with 3 µg/ml rPPE2 or TNF-α cytokine in the absence or presence of TNF-α inhibitor and the free glycerol released was quantified. (**C**) Mature 3T3-L1 adipocytes were treated with 3 µg/ml of rPPE2 and after 24 hours, cells were harvested for semi-quantitative reverse transcriptase PCR analysis of PDE3B gene (**Ci**) and densitometric quantification was carried out using ImageJ software (**Cii**). (**D**) In a separate experiment, rPPE2-treated 3T3-L1 mature adipocytes were harvested after 6 hours and 24 hours, and the levels of phosphorylated hormone-sensitive lipase (p-HSL) and total HSL were assessed by Western blotting using anti-p-HSL Ab and anti-HSL Ab respectively (**Di**). Densitometric quantification of the Western blots was carried out using ImageJ software (**Dii**). (**E**) Glycerol release was quantified in rPPE2-treated (3 µg/ml) mature 3T3-L1 adipocytes at 24 hours post-treatment in the absence or presence of H89, a PKA-specific inhibitor. (**A-E**) All the data represent mean ± SEM of 3 independent experiments. (**F**) Levels of non-esterified free fatty acid (NEFA) measured in sera of mice either left uninfected or infected with various strains of *M. tuberculosis* at day 60 post-infection. (**G**) NEFA levels in the plasma of healthy controls and TB patients. The results shown are mean ± SEM of 5 mice. For figures A, B one-way ANOVA followed by Dunnet’s and Tukey’s HSD test were carried out respectively. For figure E-F one-way ANOVA followed by Tukey’s HSD test was carried out. For figure Cii, Dii and G, unpaired Student’s *t* test was carried out. p value of < 0.05 was considered to be significant.

Since PPE2 was found to promote lipolysis, we next measured serum non-esterified free fatty acids (NEFA) levels in these mice. It was found that the mice infected with wild-type Mtb and *ppe2*-complemented CDC1551-ΔPPE2 strains had higher NEFA content compared to those infected with CDC1551-ΔPPE2 strain and uninfected control mice at 60 dpi (Figure 6F). From these data, it appears that PPE2 contributes to sustain lipolytic activity and elevated NEFA levels in circulation during mycobacterial infection. To further validate this hypothesis, we measured the plasma NEFA content of TB patients (n = 51, Supplement file 1) and compared it with those of healthy controls (n = 14). The TB patients were found to have significantly higher NEFA content compared to heathy controls (Figure 6G), suggesting presence of dyslipidaemia, an established risk factor for development of insulin resistance.

## Discussion

The association of tuberculosis and diabetes has been suspected for almost a century (*Root, 1934*). Diabetic patients are at 3-fold increased risk of developing tuberculosis compared with normoglycemic individuals (*Jeon and Murray, 2008*) and TB-patients with diabetic conditions tend to have higher treatment failures and mortality (*Dooley et al., 2009; Viswanathan et al., 2014*). Conversely, insulin resistance has been reported in TB patients without a family history of diabetes (*Magee et al., 2018; Lin et al., 2017*), while 8-87% of TB patients are newly diagnosed with diabetes (*Mave et al., 2017; Magee et al., 2018*). However, the exact mechanism that trigger development of diabetes in TB patients are not well understood.

Adipose tissue is a dynamic endocrine organ regulating glucose homeostasis and lipid metabolism (*Coelho et al., 2013*). Dysregulation of adipose tissue functions and inflammatory conditions like obesity can promote insulin resistance and type 2 diabetes. Interestingly, Mtb can persist in the adipose tissue during active TB (*Neyrolles et al., 2006; Agarwal et al., 2014*) which can modulate adipose tissue behavior (*Erol, 2008; Neyrolles et al 2006; Beigier-Bompadre et al., 2017*). Mycobacterial persistence in adipose tissue is associated with adipocyte hypertrophy, lipolysis, change in physiology and immune-cell infiltration (*Beigier-bompadre et al., 2017; Martinez et al., 2019; Bisht et al., 2023a*). These observations suggest that Mtb proteins may contribute to infection-associated hyperglycaemia, insulin resistance and metabolic dysregulation.

We previously demonstrated that PPE2, a PE/PPE family protein associated with Mtb virulence, is secreted into the circulation (*Bisht et al., 2023b*) and translocate into host nuclei (*Bisht et al., 2023b; Pal et al., 2021*). It may therefore exert systemic effects on various host tissues including adipose tissue, now recognized as a significant reservoir for Mtb (*Ayyappan et al., 2019; Neyrolles et al., 2006; Agarwal et al., 2014*). Therefore, we speculated that PPE2 is one of the potential etiological factors from Mtb that can interfere with adipocyte functions including those governing lipid metabolism, insulin sensitivity, and inflammatory responses.

To test this hypothesis, we first examined whether recombinant PPE2 protein inhibits adipogenesis in 3T3-L1 fibroblasts, since impaired adipogenesis is linked to insulin resistance and reduced abdominal fat content in mice (*Gustafson et al., 2015; Ji et al., 2014*). PPE2 protein was found to significantly inhibit adipogenesis by suppressing expression of adipogenic transcription factors PPAR-γ and C/EBP-α. PPAR-γ, regarded as “master regulator” of adipogenesis (*Farmer, 2006; Park et al., 2016*) and, together with C/EBP-α, regulates early adipogenesis events, fatty acid transport and insulin signaling/sensitization (*Wu et al., 2020; LeBlanc et al., 2016; Gerstner et al., 2024*). Mice infected with non-pathogenic *M. smegmatis* strain expressing Mtb PPE2 showed reduced adipose tissue mass accompanied by marked decrease in PPAR-γ and C/EBP-α expression. Similar changes occurred in mice administered with rPPE2 protein which could be prevented by preimmunization with rPPE2. In addition, downstream targets of PPAR-γ, such as fatty acid synthase (FAS) and adiponectin, were downregulated upon exposure to PPE2 both *in vitro* and *in vivo*. These data suggest a broad suppression of the PPAR-γ signaling axis implicating its possible involvement in tuberculosis-induced metabolic perturbations (*Oswal et al., 2022*). While the Mtb aerosol model can be questioned for bacterial load effects, it provides crucial *in vivo* validation that PPE2 function is relevant in the context of mycobacterial infection.

Weight loss and wasting are significant clinical feature associated with TB which greatly influences disease severity and prognosis. TB patients undergo significant changes in the whole-body compositions with substantial depletion of fat mass which could due to combination of several factors like less energy intake due to loss of appetite, altered metabolism due to inflammatory responses (*Paton and Ng, 2006*) and is also associated with TB-related complications like exacerbated pulmonary pathology (*Ayyappan et al., 2019*). In this study, both rPPE2 treatment and mice infection models showed significant loss in both perigonadal (PG) and visceral (VS) adipose tissue mass. In contrast, PPE2-immunized mice exhibited less severe fat loss in both tissues. Dual-energy X-ray absorptiometry (DEXA) analysis also confirmed that total body fat loss was resisted in PPE2-immunized mice. Although food intake was not quantitatively assessed, we speculate that neutralization of PPE2 by immunization was responsible for this resistance. Fat loss during TB is likely driven by chronic inflammation in the adipose tissue and production of TNF-α, IL-2, IL-1β and IFN-γ during mycobacterial infection (*Kim et al., 2011; Beigier-Bompadre et al., 2017*) some of which were also found to be upregulated in the adipose tissue transcriptome of wild-type Mtb-infected mice.

Adipocyte dysfunction often involves adipocyte hypertrophy, increased oxidative stress, chronic low-grade inflammation and impaired adipocytes from its precursor pre-adipocytes (*Klöting and Blüher, 2014; Reyes-Farias et al., 2021*). Under normal physiological conditions, the major roles of adipose tissue include insulin-stimulated glucose uptake, storage of excess energy as triglyceride, and endocrine functions like secretion leptin, resistin and cytokines (*Scheja and Heeren, 2019*). Impaired adipogenesis is associated with insulin resistance (*Hammarstedt et al., 2018*), while adipocyte hypertrophy promotes immune cell infiltration, oxidative stress, NLRP3 inflammasome activation leading to production of pro-inflammatory cytokines which contribute to development of insulin resistance (*Kursawe et al., 2016; Susca et al., 2024*).

Interestingly, mice infected with *ppe2-*null mutant showed modest hypertrophy, despite a lower bacterial burden and absence of PPE2. This suggests that hypertrophy can be attributable to additional PPE2-independent, infection-associated mechanisms. Mycobacterial cell envelope components like lipoproteins, peptidoglycans, lipoarabinomannans may trigger pattern-recognition receptor-mediated signaling through TLR2/TLR4 and inflammatory responses (e.g., TNF-α, IL-6), which are known to promote adipocyte hypertrophy and lipid accumulation. Additionally, metabolic demand for mounting an immune response against mycobacterial infection, may redirect host energy and lipid homeostasis. This is consistent with our earlier observation in *M. smegmatis* infection model, where a vector-control *M. smegmatis* strain also resulted in a modest reduction in adipose mass. Thus, PPE2 appears to exacerbate rather than exclusively, cause infection-associated adipose tissue remodelling. Therefore, we speculate that PPE2-independent mycobacterial factors also contribute to modest adipose tissue remodelling during Mtb infection.

We also observed that PPE2 can promote immune cell infiltration in adipocytes, including T-cells and macrophages. Immune cell infiltration is widely implicated as an important factor for adipose tissue dysfunction and development of insulin resistance (*Guzik et al., 2017*). Infiltration of T-cells and macrophages into perigonadal adipose tissue, as also observed in our study, was found to be associated with altered gene expression and elevated interferon-γ signaling causing localized tissue inflammation (*Beigier-Bompadre et al., 2017*). Inflammation can promote lipolysis, leading to elevated circulating free fatty acids, and thereby contributing to peripheral insulin resistance (*Beigier-Bompadre et al., 2017; Guilherme et al., 2008*). Furthermore, PPE2 also inhibited adiponectin expression followed by administration of rPPE2 or infection with *M. smegmatis* expressing PPE2. Adiponectin has been implicated to play an important role in insulin sensitivity and glucose homeostasis. Therefore, inhibition of adiponectin expression by PPE2 appears to be another mechanism by which it can promote glucose intolerance and subsequent development of insulin resistance.

A role of PPE2 in modulation of adipose tissue physiology was further confirmed by using a *ppe2*-null mutant (CDC1551-ΔPPE2) of a clinical *M. tuberculosis* (CDC1551) strain. Suppression of PPAR-γ and CD36 expression was alleviated in CDC1551-ΔPPE2-infected mice, whereas genetic complementation restored wild-type-like phenotype. Functionally complemented (CDC1551-ΔPPE2::PPE2) and wild-type strains showed exacerbated adipose tissue pathology compared to those infected with CDC1551-ΔPPE2 strain which showed almost normal adipose tissue histology comparable to uninfected controls At 60 days post-infection, mice infected with wild-type as well as CDC1551-ΔPPE2::PPE2 showed pronounced insulin resistance as assessed by HOMA-IR. At 60 days post-infection, A noticeable decrease in PG and VS fat tissue weight was observed in all mice as compared to day 30 post-infection which could be due to age-related fat mass atrophy.

To the best of our knowledge, our data provide first time evidence that Mtb PPE2 is one of the possible etiological factors that contribute to adipocyte dysfunction, metabolic abnormalities relevant to TB-associated hyperglycemia and insulin resistance. PPE2 was found to suppress PPAR-γ and C/EBP-α, promotes adipocyte hypertrophy, reduces adiponectin expression, promotes immune cell infiltration and lipolysis. These processes possible orchestrate together to impair glucose homeostasis. PPE2 also altered the adipose tissue transcriptome, affecting genes involved in wide ranges of physiological functions, which include tissue-specific immune responses, phospholipase C activity, ribosomal biogenesis and pathways involved in neurodegenerative disorders. Such broad effects are in concordance with our earlier observation that PPE2 possesses features of a eukaryotic transcription factor including a functional nuclear localization signal (NLS) and a DNA-binding domain, enabling nuclear translocation and binding to GATA-1-like elements (*Bhat et al., 2017; Pal et al., 2021*). PPE2 also contains a SH3 motif (*Srivastava et al., 2019*), a protein-protein interaction domain, characteristic of several kinases involved in cellular signaling (*Kurochkina and Guha, 2013*). These properties likely to have a profound influence on the transcriptome as well as the protein-protein interactome of the adipose tissue. The data presented herein clearly demonstrated a definitive role of PPE2 in modulating adipose tissue physiology which is not merely due to differences in bacterial burden.

Although we did not explore the precise mechanisms by which PPE2 regulate adipogenic transcription factors and adiponectin, our findings suggest possible mechanisms by which PPE2 promote lipolysis which can trigger development of insulin resistance by promoting inflammation (*Morigny et al., 2016*). PPE2 was found to trigger ER stress in the adipose tissue, which in turn activate cAMP/PKA causing increased activation of HSL leading to increased lipolysis of adipocyte triglycerides. Increased lipolysis may elevate circulating NEFA, and consistent to this hypothesis, serum NEFA was found to be significantly higher in active TB patients. However, this clinical observation does not establish a direct role of PPE2 in human context and studies correlating circulating PPE2 levels with serum NEFA and metabolic parameters in a well-characterized TB patient cohort will be needed. The observation that mice infected with wild-type Mtb or the complemented strain CDC1551-ΔPPE2::PPE2 exhibited elevated serum NEFA levels compared to those infected with *ppe2*-deficient CDC1551-ΔPPE2 strain suggests that PPE2 probably contributes to increased circulating NEFA levels during mycobacterial infection. Since, pre-immunization with rPPE2 protein was able to counter the adverse effects of PPE2 in mouse models, it will be interesting to further investigate whether PPE2-based subunit vaccine can effectively mitigate TB-associated hyperglycaemia and insulin resistance.

## Materials and methods

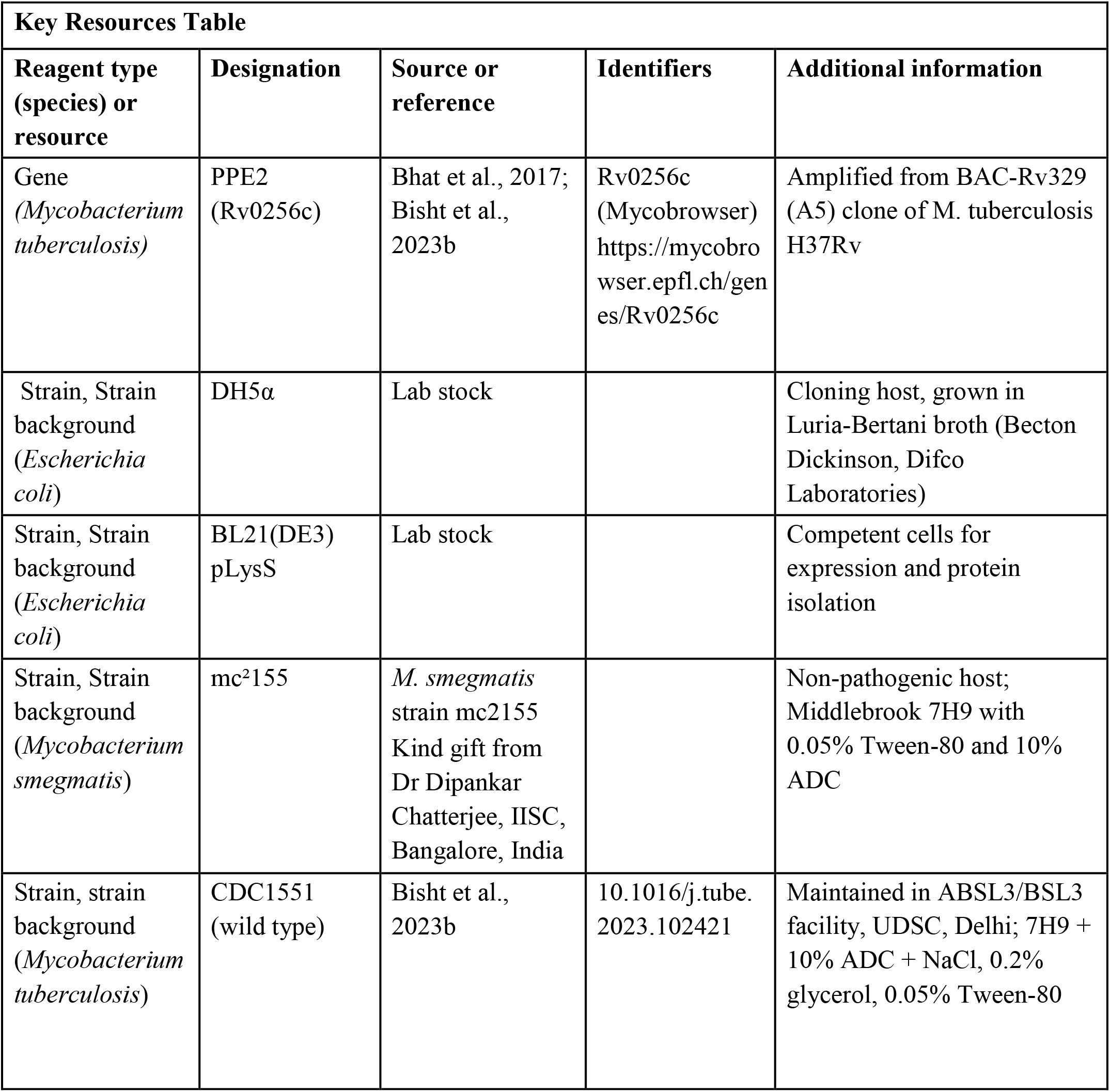
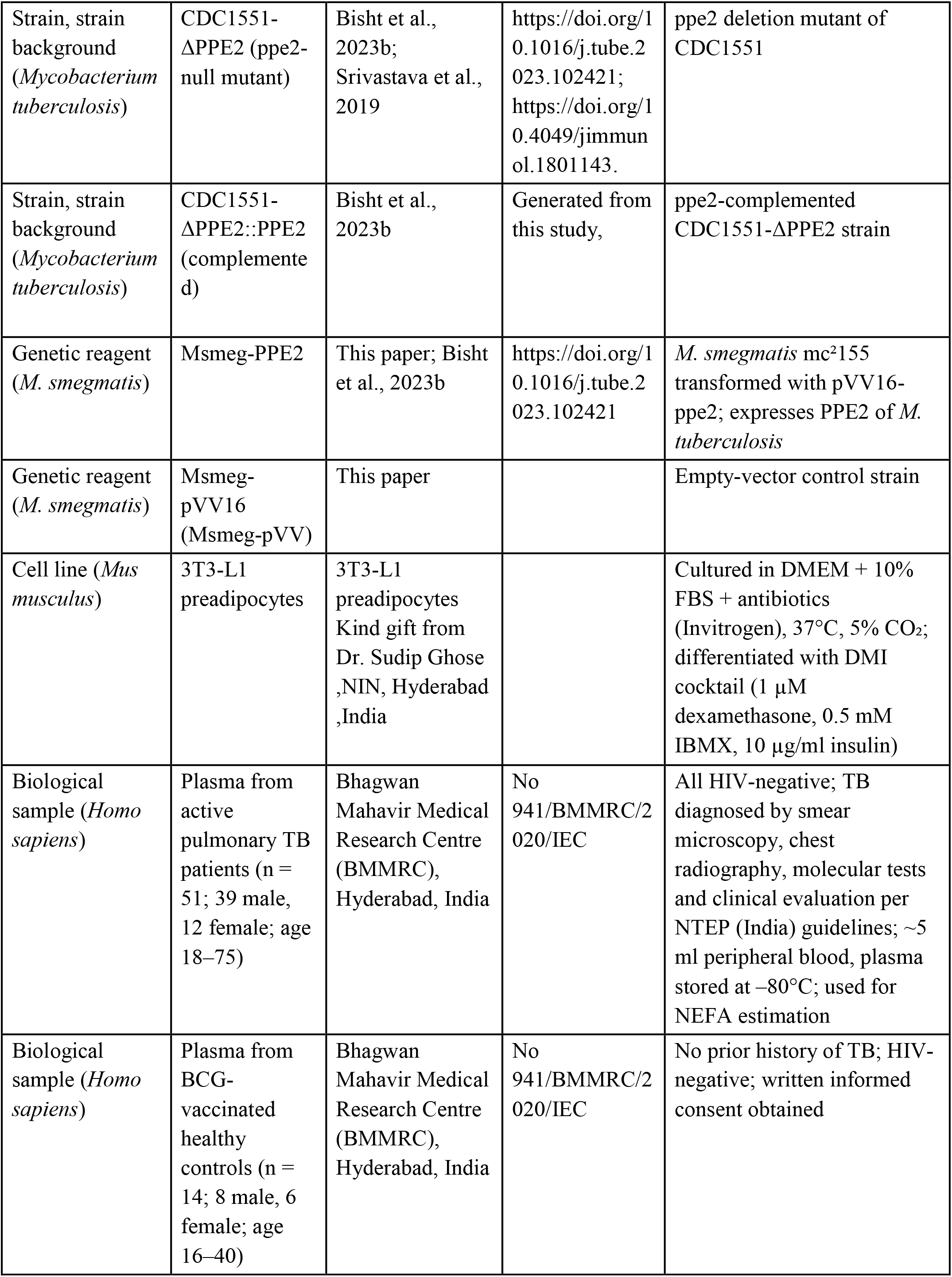
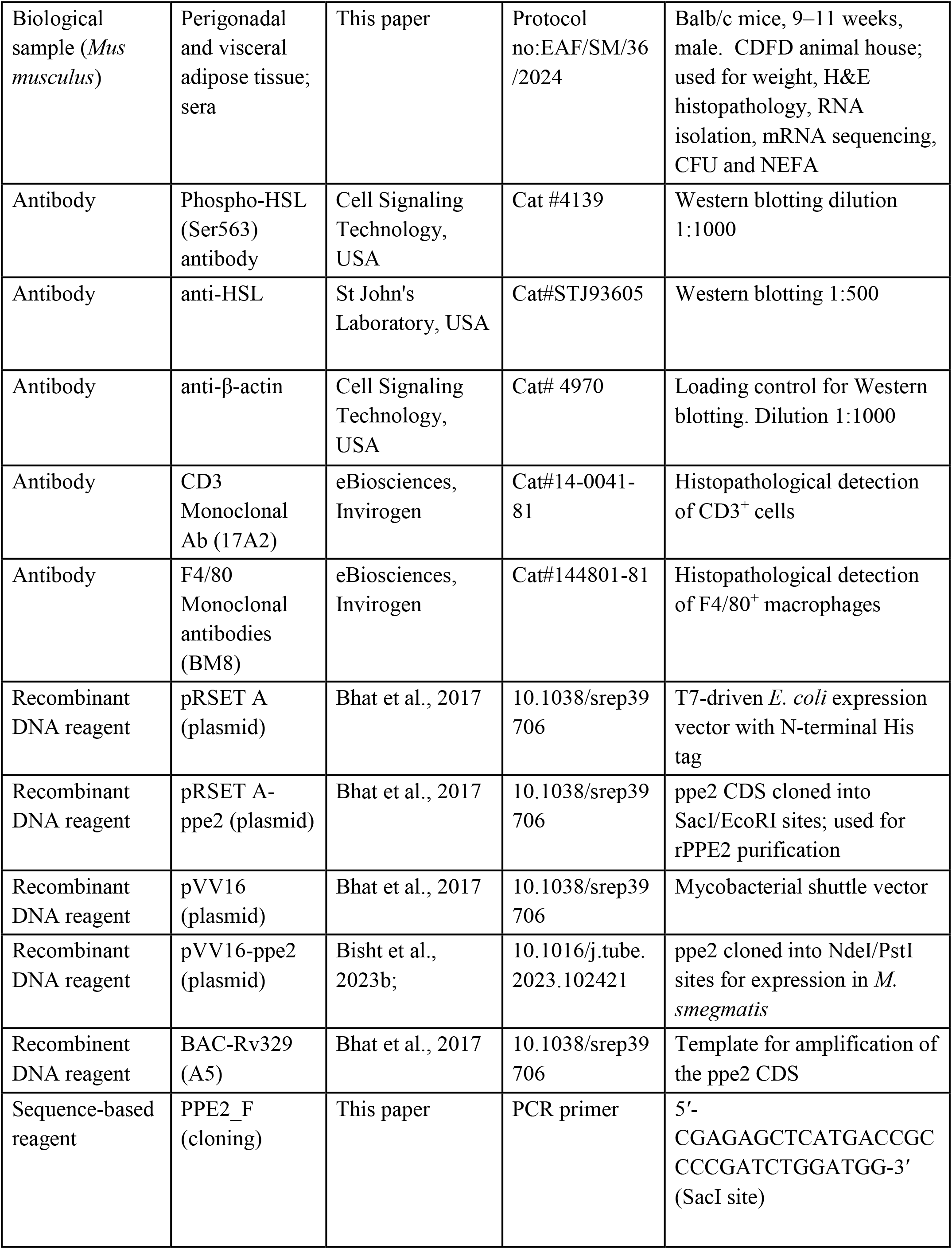
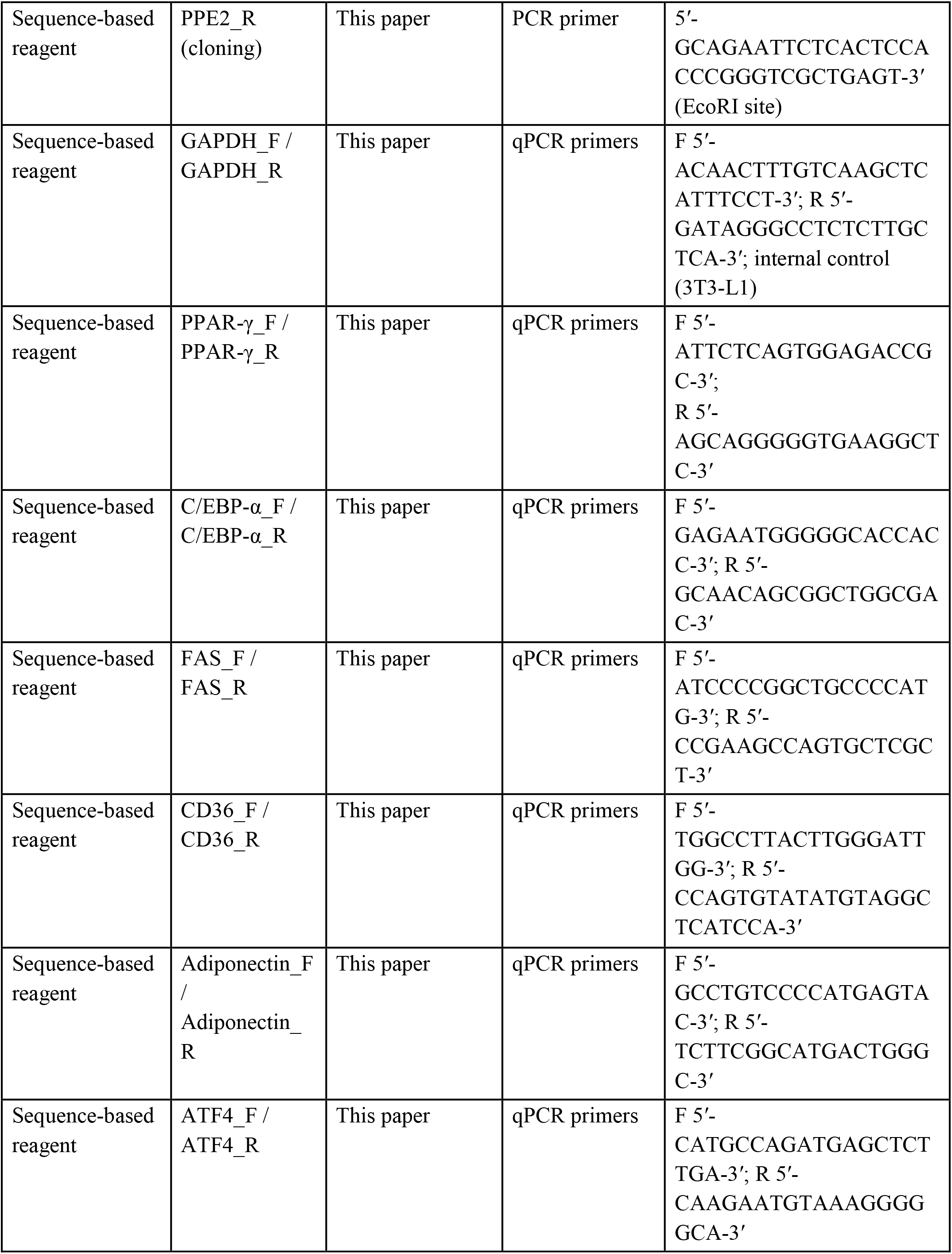
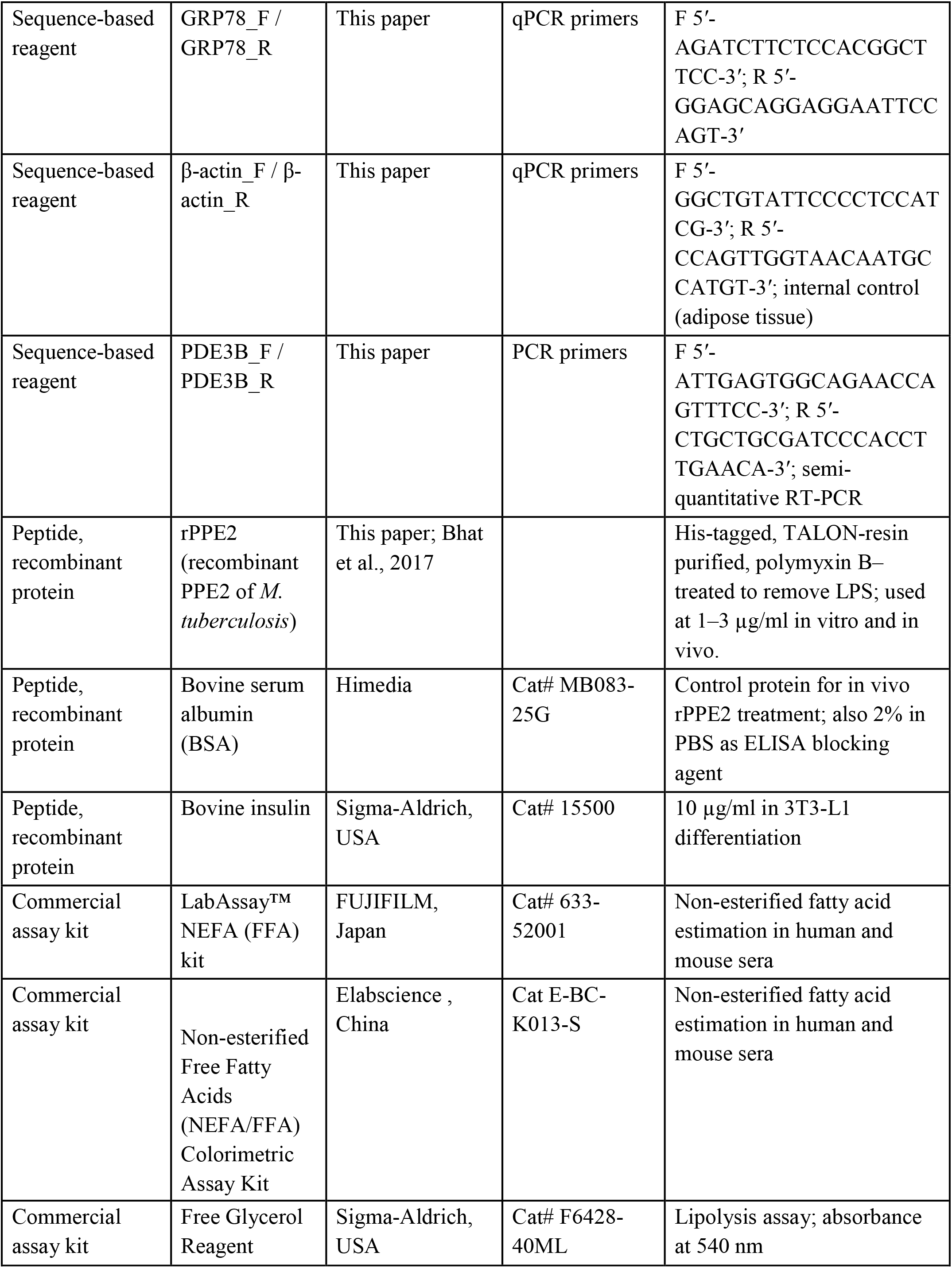
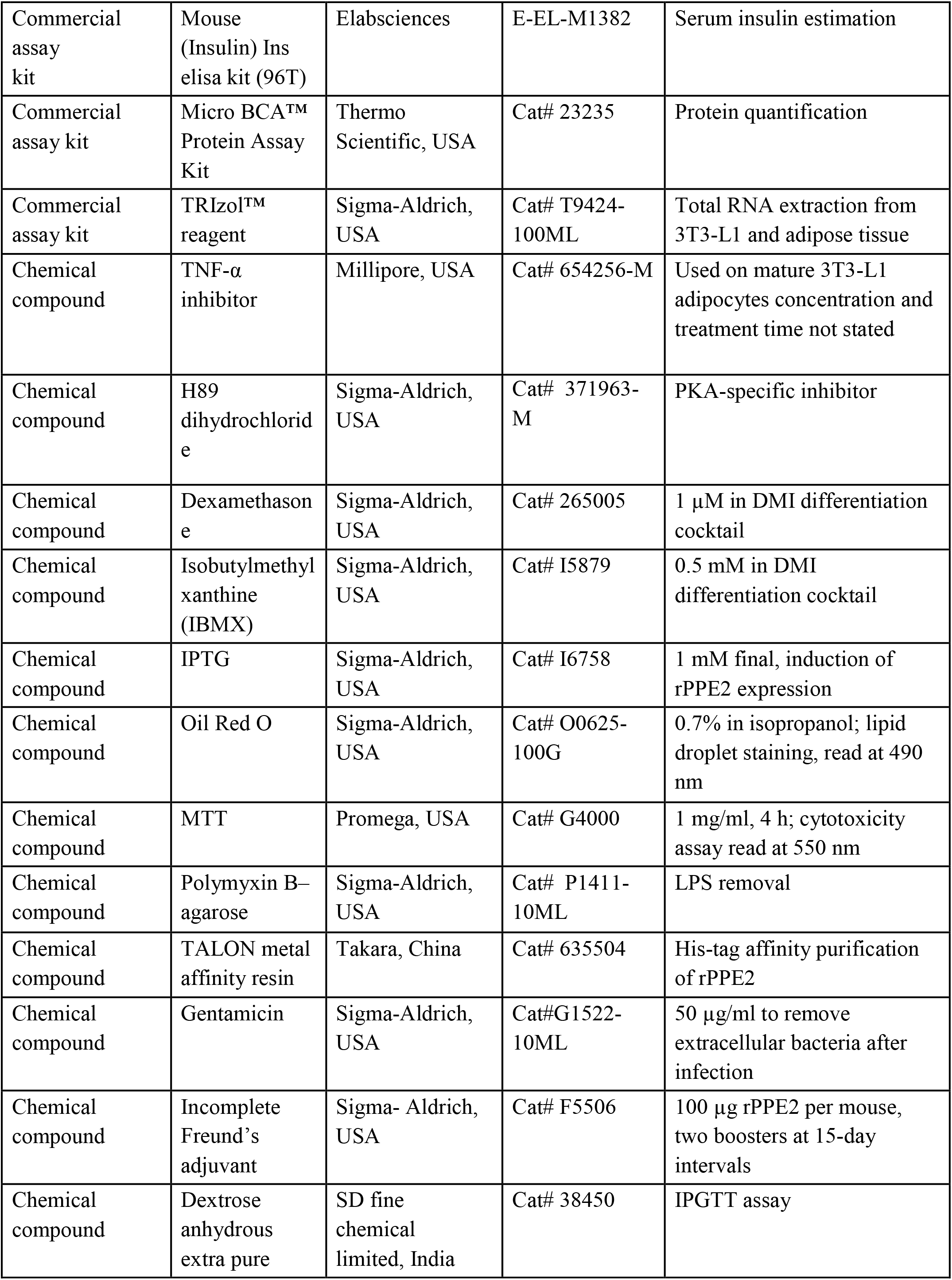
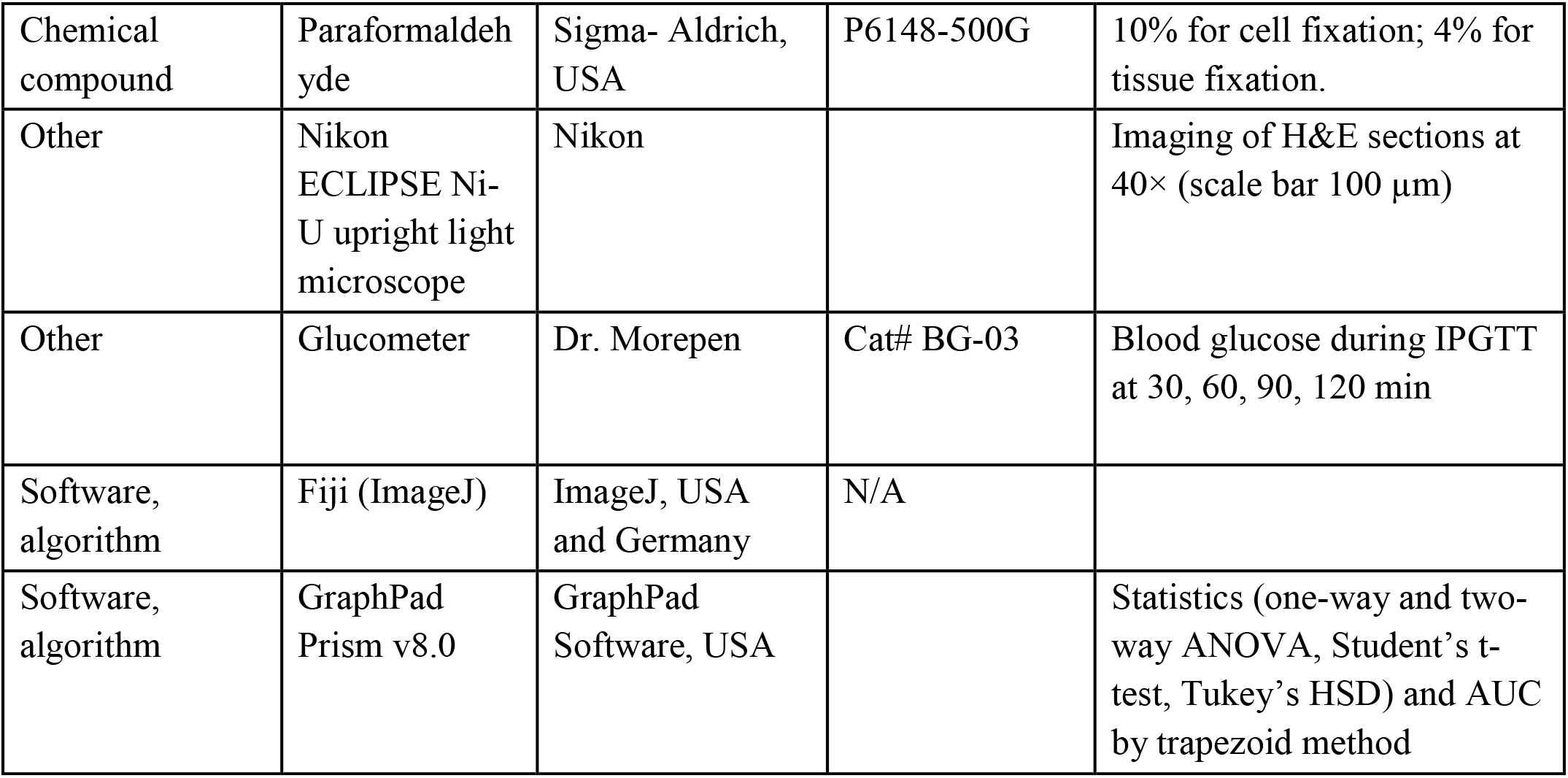

### Ethical statement and sample collection

All animal experimental protocols were approved by the Institutional Animal Ethics Committee of BRIC-Centre for DNA Fingerprinting and Diagnostics (BRIC-CDFD), Hyderabad, India. Balb/c mice were maintained at the animal house facility of CDFD, Hyderabad, under standard conditions. The study involving human sample collection was approved by the Institutional Ethical Committee for Biomedical Research, at Bhagwan Mahavir Medical Research Centre (BMMRC) (No 941/BMMRC/2020/IEC). The study included 51 active pulmonary TB patients and 14 BCG-vaccinated healthy controls with no prior TB history. About 5 ml peripheral blood was collected, plasma was separated and stored at –80 °C until further use. The control group (8 males, 6 females; age 16–40) and TB group (39 males, 12 females; age 18–75) were all HIV-negative. All the samples were collected after obtaining written informed consents from the study participants. Active TB was diagnosed using smear microscopy, chest radiography, molecular tests, and clinical evaluation, following NTEP guidelines (India).

### Reagents

Antibodies and reagents used in this study are anti-phospho-HSL (Cell Signaling Technology, USA), anti-HSL antibody (St Jhon Laboratory, USA), anti-β-actin antibody (Cell Signaling Technology, USA), TNA-α inhibitor (654256-M, Millipore, USA), H89 dihydrochloride (H89, Santa Cruz, USA), NEFA Kit (FUJIFILM, Japan). Free glycerol reagent was purchased from Sigma-Aldrich, USA (F6428-40mL).

### Purification of recombinant PPE2 protein (rPPE2)

PPE2 protein was purified as described by us earlier (*Bhat et al., 2017*). For affinity purification of the recombinant PPE2 protein, the CDS was amplified from BAC-Rv329 (A5) using a forward primer (5′-CGAGAGCTCATGACCGCCCCGATCTGGATGG-3′) and a reverse primer (5′-GCAGAATTCTCACTCCACCCGGGTCGCTGAGT-3′) and cloned into the *Sac*I and *Eco*R1 site of a T7 polymerase driven *Escherichia coli* expression vector pRSET A (Invitrogen, Carlsbad, USA) in frame with an N-terminal histidine tag to facilitate purification using metal affinity purification resins. The pRSET A vector containing the PPE2 gene was then transformed into *Escherichia coli* BL21 (DE3) pLysS cells and a primary inoculum was prepared from a single transformed colony. The primary culture was inoculated into 800 ml terrific broth in the presence of chloramphenicol (35 μg/ml) and ampicillin (100 μg/ml). The culture was grown in a shaker incubator at 37°C till the absorbance reached to about 0.5. At this stage, expression of the recombinant protein was induced by adding IPTG (Isopropyl β-d-1-thiogalactopyranoside, Sigma-Aldrich, USA) to a final concentration of 1 mM and incubation was continued further for 16 hours at 18°C in a shaker incubator to favor soluble expression of the protein. Cells were then harvested by centrifugation and the pellets were suspended in lysis buffer (PBS containing 5% glycerol, 0.3% sodium lauroyl sarcosine). The cell suspension was then sonicated and the lysate was centrifuged at 12000 rpm for 30 minutes. The soluble recombinant protein present in the supernatant was allowed to bind with the TALON resin (Clontech, USA) for an hour, loaded into a column and washed with washing buffer (PBS containing 5% glycerol and 20 mM Imidazole). The resin-bound protein was eluted using elution buffer (PBS containing 5% glycerol and 200 mM imidazole). Eluted samples were loaded onto an SDS-PAGE gel to confirm the presence of the recombinant PPE2 protein. The PPE2-positive fractions were pooled together and dialyzed against PBS containing 5% glycerol to remove imidazole from the protein samples. The concentration of the purified protein was estimated using MicroBCA^TM^ Protein Assay Kit from Thermo Scientific (Rockford, USA) following manufacturer’s instructions. To remove LPS contamination, the recombinant proteins were treated with 10% (v/v) polymyxin B-agarose (Sigma-Aldrich, USA; binding capacity 200 – 500 µg of LPS/ml) for 1 hour at 4°C as described by us earlier (*Bhat et al., 2013*).

### Cloning and expression of *ppe2* gene of *M. tuberculosis* in *M. smegmatis* bacteria

The non-pathogenic *M. smegmatis* mc^2^155 bacteria were received from ATCC and were grown in Middlebrook 7H9 medium (HiMedia Laboratories, India) supplemented with 0.05% Tween-80 (Sigma-Aldrich, USA) and 10% Albumin-dextrose-catalase (ADC) (HiMedia Laboratories, India), For generating *M. smegmatis* expressing PPE2 protein of *M. tuberculosis* (Msmeg-PPE2), *ppe2* was cloned in a mycobacterial shuttle vector pVV16 into *Nde* I – *Pst* I sites for expression in *M. smegmatis* as described earlier (*Bisht et al., 2023b*). Transformation of pVV16-*ppe2* construct into *M. smegmatis* (Msmeg-PPE2) was carried out as described earlier (*Bhat et al., 2017*). *M. smegmatis* harboring the empty vector (Msmeg-pVV16) alone was used as control.

### Culture of bacterial strains

The DH5α strain of *E. coli*, used for cloning was grown in Luria-Bertani broth (Becton Dickinson, Difco Laboratories). Wild-type *M. tuberculosis* strain (CDC1551) and *ppe2-*null mutant *M. tuberculosis* strain (CDC1551-ΔPPE2) and complemented strain (CDC1551-ΔPPE2::PPE2) were maintained as described earlier (*Bisht et al., 2023b*) in ABSL3/BSL3 facility of the University of Delhi South Campus (UDSC), Delhi, India. Mycobacterial strains were grown in Middlebrook 7H9 Broth supplemented with 10% albumin, dextrose, catalase, and NaCl (HiMedia Laboratories) along with 0.2% glycerol (Sigma-Aldrich, USA) and 0.05% Tween-80 (Sigma-Aldrich, USA).

### Differentiation of 3T3-L1 preadipocytes into mature adipocyte cells

3T3-L1 fibroblast cells were differentiated into adipocytes using a differentiation cocktail containing dexamethasone, insulin and IBMX. In brief, 3T3-L1 cells (obtained from ATCC) were cultured in Dulbecco’s modified Eagle’s medium (DMEM) supplemented with 10% fetal bovine serum and antibiotics (all from Invitrogen USA) and were maintained in an incubator at 37°C and 5% of CO2. At confluence, preadipocytes were induced to differentiate into adipocytes by culturing them in differentiation cocktail (DMI) comprising of 1 μM dexamethasone (Sigma-Aldrich, USA), 0.5 mM isobutylmethylxantine (Sigma-Aldrich, USA) and 10 μg/ml bovine insulin (Sigma-Aldrich, USA). On day 2, the medium was replaced with DMEM with 10% fetal bovine serum and insulin (10 µg/ml). From day 4 onwards, medium was replaced with 10% FBS in DMEM and antibiotics but without insulin, and this medium was changed every 2 days up to day 8 after confluence, when ∼95% of the cells were differentiated to adipocytes.

### Measurement of degree of differentiation of 3T3-L1 cells in the presence of rPPE2

The degree of differentiation was measured by staining with Oil Red O as described by *Wu et al*. (2007). 3T3-L1 cells were differentiated either in the absence or presence of different concentrations of recombinantly purified PPE2 protein of *M. tuberculosis* (rPPE2). On day 10, the cells were fixed with 10% paraformaldehyde for one hour and subsequently washed with distilled water. The fixed cells were stained with 0.7% solution of Oil Red O stain (Sigma-Aldrich, USA) in isopropanol, followed by multiple washes with distilled water. To quantify Oil Red O stain retained by the lipid droplets present in matured 3T3-L1 cells, the dye was extracted using isopropanol and the absorbance was measured at 490 nm using a spectrophotometer. Also, total RNA was isolated from differentiating 3T3-L1 adipocyte culture medium containing rPPE2 (3 µg/ml) at various time points and adipogenic markers like PPAR-γ, C/EBP-α, fatty acid synthase (FAS) expression was measured by reverse transcription polymerase chain reaction (RT-PCR) using gene-specific primers.

### Effect of PPE2 on adipose tissue physiology *in vivo*

Balb/c mice of 9-11 weeks of age were infected with 100 million of either *M. smegmatis* carrying empty vector pVV16 (Msmeg-pVV) or *M. smegmatis* expressing PPE2 (Msmeg-PPE2) through intravenous route and were sacrificed on 7^th^ day post-infection. The perigonadal and visceral fat tissue were harvested for measuring the weight and also used for histopathology and extraction of total RNA. Histopathology was carried out by hematoxylin and eosin (H&E) staining.

### RNA isolation and RT-PCR and semi-quantitative reverse transcriptase PCR

Total RNA was extracted from 3T3-L1 or perigonadal and visceral fat tissue using trizol following the manufacturer’s protocol (Invitrogen, USA). After converting the total RNA into cDNA, the levels of expression of PPAR-γ, C/EBP-α, FAS, CD36 and Adiponectin was measured by qPCR. GAPDH was used as an internal control. The primers used are as follows; GAPDH (F 5’ACAACTTTGTCAAGCTCATTTCCT3’ and R 5’GATAGGGCCTCTCTTGCTCA3’), PPAR-γ (F 5’ATTCTCAGTGGAGACCGC3’ and 5’AGCAGGGGGTGAAGGCTC3’), C/EBP-α (F 5’GAGAATGGGGGCACCACC3’ and R 5’GCAACAGCGGCTGGCGAC3’), FAS (F 5’ATCCCCGGCTGCCCCATG3’ and R 5’CCGAAGCCAGTGCTCGCT3’), CD36 (F 5’ TGGCCTTACTTGGGATTGG3’ and R 5’CCAGTGTATATGTAGGCTCATCCA3’) and Adiponectin (F 5’GCCTGTCCCCATGAGTAC3’ and R 5’TCTTCGGCATGACTGGGC3’). Also, the levels of expression of ATF4 (F 5’CATGCCAGATGAGCTCTTGA3’ and R (5’CAAGAATGTAAAGGGGGCA3’) and GRP78 (F 5’AGATCTTCTCCACGGCTTCC3’ and R (5’GGAGCAGGAGGAATTCCAGT3’) in perigonadal fat tissue were checked where β-actin (F 5’GGCTGTATTCCCCTCCATCG 3’ and R 5’CCAGTTGGTAACAATGCCATGT3’) was used as internal control. PDE3B level in 3T3-L1 was measured by semi-quantitative reverse transcriptase PCR using the following primer; PDE3B (F 5’ ATTGAGTGGCAGAACCAGTTTCC3’ and R 5’ CTGCTGCGATCCCACCTTGAACA3’).

### Protein treatment in mice

Balb/c mice of 9-11 weeks old mice were administered with either bovine serum albumin (BSA) or rPPE2 intravenously on alternate days for 12 days. On day 14, glucose tolerance test was performed using standard protocol as described in the following section. Mice were sacrificed on day 15^th^ and perigonadal and visceral adipose tissue weights were measured. Also, perigonadal and visceral tissues were harvested for histopathology analyses using H&E staining and RNA isolation was carried out for measuring the expression of PPAR-γ, C/EBP-α and Adiponectin.

### Histopathology

For histopathology analyses, tissue samples were fixed with 4% paraformaldehyde and embedded in paraffin wax, and then sectioned (4∼5 μm) for hematoxylin and eosin (H&E) staining. For this, the sections were deparaffinized and hydrated in distilled water and stained with Hematoxylin (30 seconds), washed in distilled water, and counterstained with Eosin (1% in distilled water for 30 seconds). Slides were washed, dehydrated, and mounted on slides for imaging. Tissue section images were acquired using Nikon ECLIPSE Ni-U light upright microscope.

### Intraperitoneal glucose tolerance test (IPGTT)

Balb/c mice were starved overnight and an intraperitoneal injection of glucose was given at a dose of 2 g/kg as 25% glucose solution in sterile saline. Blood glucose levels were monitored from the tail tip vein using a glucometer at 30, 60, 90 and 120 minutes post-injection. The data plotted as percent of initial glucose concentration *versus* time (*Veloso et al., 2019; Martinez et al., 2019*). The area under the curve was calculated using the trapezoid method using Graph Pad Prism (GraphPad Software, USA).

### *M. tuberculosis* infection

Balb/c mice of 9-11 weeks of age were aerosol-infected with approximately 200-300 CFU per mice of either CDC1551 (wild-type *M. tuberculosis) or CDC1551-ΔPPE2 *or* CDC1551-ΔPPE2::PPE2 (ppe2-*complemented CDC1551-ΔPPE2 strain) and sacrificed at 30 days post-infection (dpi) and 60 dpi respectively. The perigonadal and visceral fat tissue were harvested and the weight of these tissues was measured. The histopathology of perigonadal and visceral fat tissue was performed using hematoxylin and eosin (H&E) staining. Additionally, RNA was extracted from perigonadal and visceral fat tissue using Trizol following the manufacturer’s protocol (Invitrogen, USA) and the levels of expression of PPAR-γ and CD36 were measured by qPCR. Perigonadal adipose tissue from mice sacrificed at day 60 post-infection was used for mRNA sequencing study.

### Transcriptome analysis

Total RNA was extracted from the perigonadal fat tissues from uninfected, wild-type- and CDC1551-ΔPPE2-infected mice using Direct-ZOL Mini Prep Kit (Zymo Research) as per the manufacturer’s protocol. RNA quality and quantity were assessed using a NanoDrop spectrophotometer and Agilent Tape station with high-sensitivity RNA ScreenTape. Paired-end RNA-sequencing libraries were prepared using NEBNext^®^ Ultra^TM^ II Directional RNA Library Prep Kit for Illumina (NEB) as per the manufacturer’s protocol. The libraries were purified using AMPureXP beads and analyzed on 4200 Tape Station system (Agilent Technologies) using high-sensitivity D1000 Screen tape as per the manufacturer’s protocol. The sequencing was performed on an Illumina MiSeq platform. The raw reads were processed using Trimmomatic v0.39 to remove adaptor sequences, ambiguous reads (reads with unknown nucleotide ‘N’ larger than 5%), and low-quality sequences (reads with more than 10% quality threshold (QV) < 25phred score). High-quality reads with a minimum length of 100 nucleotides (nt) after trimming were retained. These high-quality reads (QV>25) were mapped on the mouse reference genome (GRCm39.110) obtained from ENSEMBLE using STAR (v2.7.10a) using default parameters. FeatureCounts (version 2.0.3) was used to count the number of reads mapped to each gene. Differential gene expression analysis was performed using the DESeq2 R package between the groups. Log2 Fold Change values greater than zero were considered upregulated, while those less than zero were downregulated, along with a P-value threshold of 0.05 for statistically significant results. Finally, the gene set enrichment analysis (GSEA) was performed using software GSEA (4.3.3) from the UC San Diego and Broad Institute website following the standard data processing pipeline (GSEA (gsea-msigdb.org)) (*Subramanian et al., 2005; Mootha et al., 2003*).

### Colony forming unit (CFU) count

For CFU count, perigonadal adipose tissue from *M. tuberculosis*-infected mice was homogenized in saline solution and plated on 7H11 plates supplemented with 10% OADC (HIMEDIA, India). Plates were next incubated at 37°C and the colonies were counted after 30-35 days post-plating.

### NEFA estimation

Non-esterified free fatty acids (NEFA) were analyzed in human and mouse sera samples using LabAssay™ NEFA (FFA) kit from FUJIFILM, Japan, following the manufacturer’s protocols.

### Western blotting

3T3-L1 cells were lysed using RIPA buffer (50 mM Tris-HCl [pH 8], 150 mM NaCl, 2 mM EDTA, 1% NP-40, 0.5% Sodiumdeoxycholate, 0.1% SDS and protease inhibitors) and the proteins were resolved on SDS-PAGE followed by Western blotting using specific primary antibody and appropriate combinations of HRP (horseradish peroxidase) conjugated secondary Ab (Sigma-Aldrich, USA). Bound HRP enzyme was detected by chemiluminescence following the manufacturer’s protocol (GE Healthcare, UK).

### Measurement of lipolysis

Lipolysis in mature 3T3-L1 adipocytes was measured by estimating free glycerol released in the medium using Free Glycerol Reagent Kit (F6428-40mL, Sigma-Aldrich, USA) following the manufacturer’s protocol. Briefly, 10 µl of culture medium was added to 800 µl of free glycerol reagent and kept at 37℃ for 5 minutes. The reaction results in production of a quinoneimine dye which was measured at 540 nm using a spectrophotometer. The results were expressed as µg of glycerol released per mg protein.

### Statistical analysis

GraphPad Prism software, version 8.0 was used for determining the significance of the difference in the mean between the samples. One-way ANOVA, two-way ANOVA and Student’s t-test was carried out followed by appropriate post-hoc test to determine the significance of difference. A p value of < 0.05 was considered significant.

## Acknowledgement

We thank UDSC-ABSL3/BSL3 facility for providing access to the BSL3 facility at the University of Delhi South Campus for the work related to the use of the pathogenic strain of *Mycobacterium tuberculosis.* We thank Dr Garima Khare from UDSC and her team members, Dr Prachi Nagpal and Dr Nupur Angrish, for their assistance during the mouse aerosol infection. We also thank Dr Sandeep Kumar Kotturu and Dr. N.V. Sanjeev Kumar for kind help. The authors gratefully acknowledge the financial support by the Science and Engineering Research Board (SERB), Department of Science and Technology (DST), Government of India (JCB/2021/000035), Council of Scientific and Industrial Research (CSIR), Govt. of India (37WS(0020)/2023-24/EMR-II/ASPIRE), Department of Biotechnology (DBT), Govt of India (BT/PR51149/MED/29/1660/2023) to SM and SG, Indian Council of Medical Research (ICMR), Govt. of India (2021-10087/GTGE/ADHOC-BMS and IIRPSG-2024-01-01453) and a core grant from CDFD by DBT. VM was supported by the DST-INSPIRE fellowship, Govt. of India.

## Declaration of interests

A patent application based on the results described in this paper is being filed by CDFD, Hyderabad, in which S.M. M.K.B and S.G. are listed as inventors.

## Author contributions

M.K.B., S.G. and S.M. conceived, designed the experiments, and analyzed the data. M.K.B. performed all the experiments. V.M. and P.D. helped in 3T3-L1 and mouse infection experiments. V.L.V. provided healthy control and TB patients’ plasma samples. M.K.B., S.G. and S.M. wrote the manuscript. S.G. and S.M. corrected and edited the manuscript.

## Supplementary Figures

**Figure 1—figure supplement 1.**
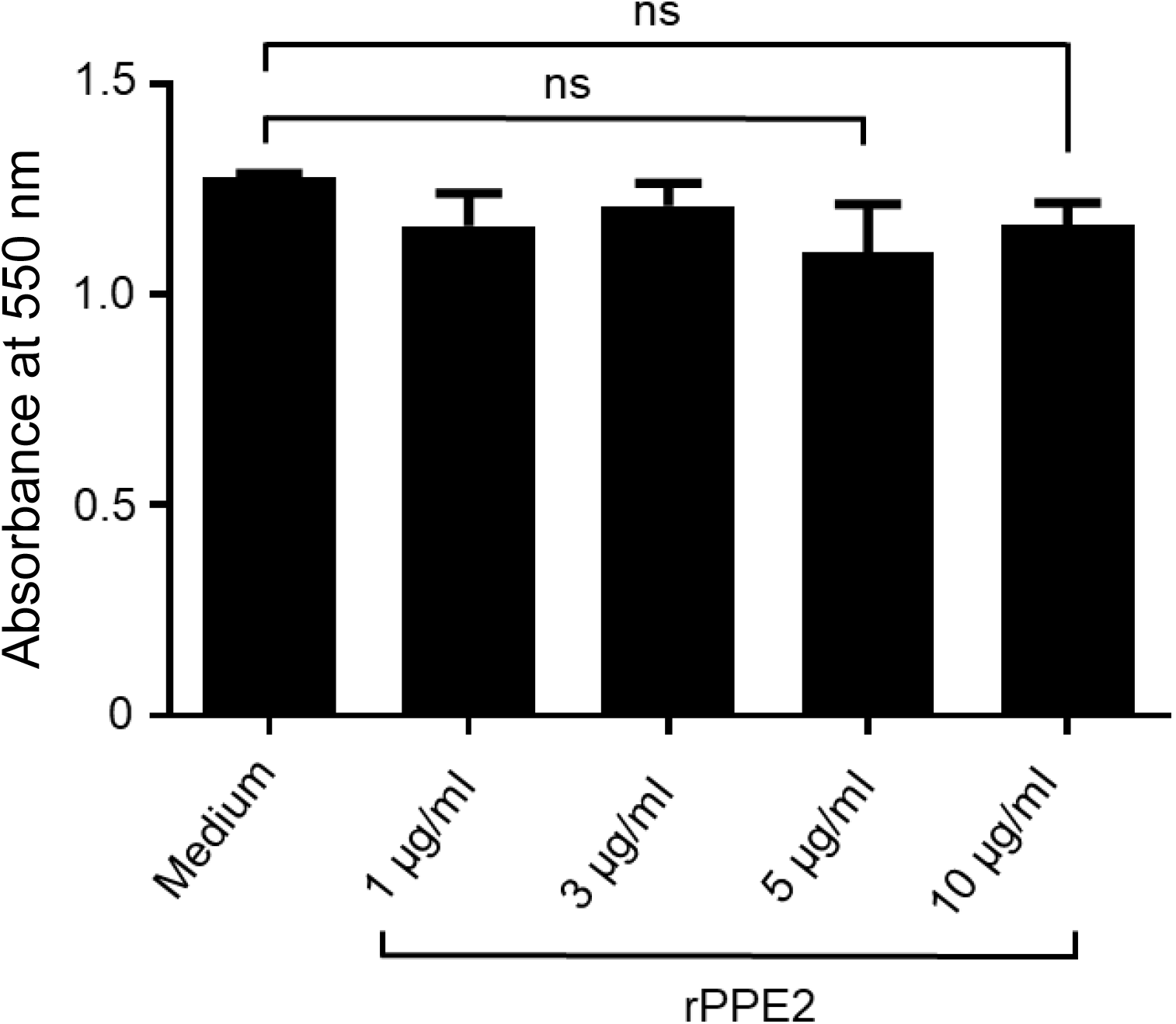
rPPE2 does not cause 3T3-L1 cell toxicity. 3T3-L1 cells (2 × 10^5^/ml) were treated with different concentrations of rPPE2 for 48 hours and cell cytotoxicity was measured by MTT assay. In brief, MTT (3-(4,5-Dimethylthiazol-2-yl)-2,5-diphenyltetrazolium bromide, Sigma-Aldrich, USA) was added at 1 mg/ml and incubated for 4 hours. The cells were lysed overnight using 100 μl of lysis buffer (20% SDS and 50% dimethyl formamide) and the absorbance was measured at 550 nm. One-way ANOVA followed by Tukey’s HSD test was carried out. ns = non-significant, p-value of < 0.05 was considered to be significant.

**Figure 1—figure supplement 2.**
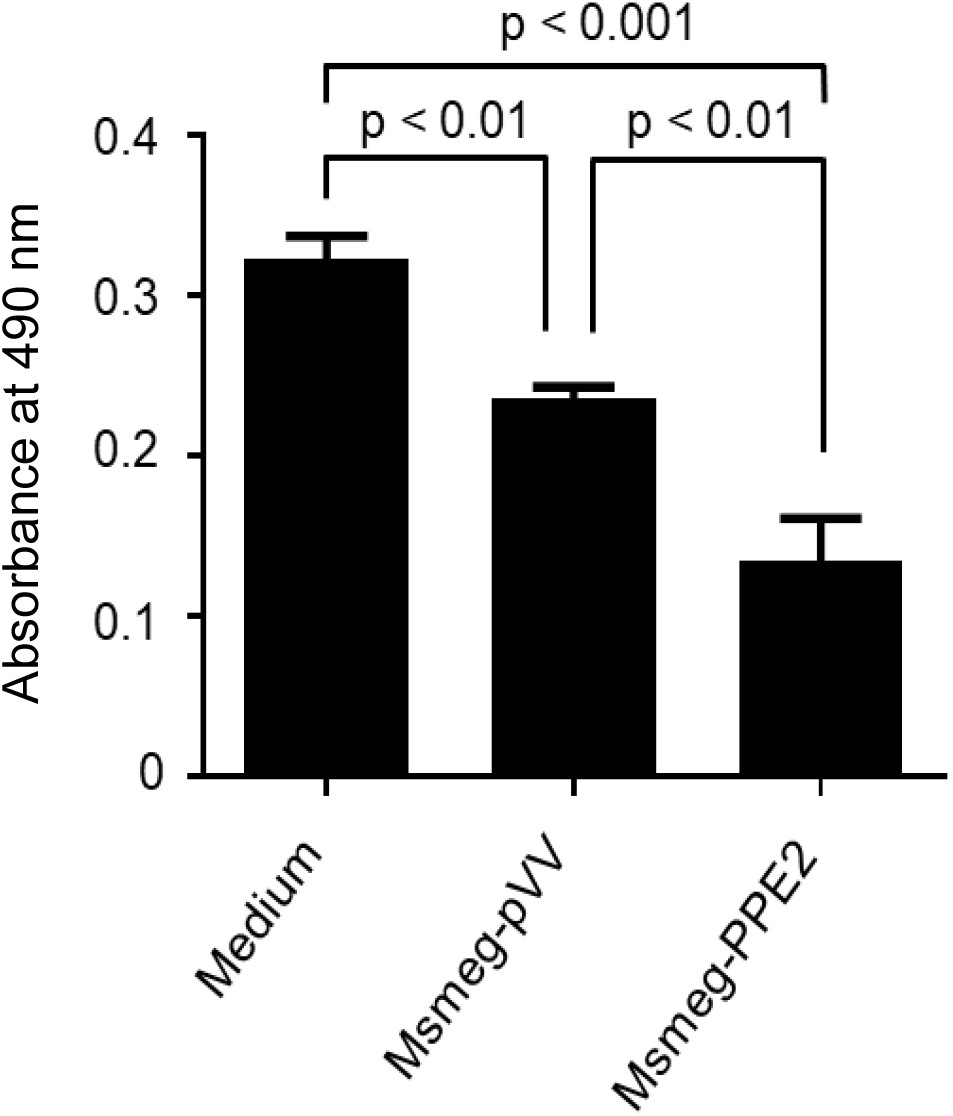
PPE2 inhibits 3T3-L1 adipocyte differentiation during infection. Undifferentiated 3T3-L1 pre-adipocytes were infected with *M. smegmatis* harboring pVV16 empty vector (Msmeg-pVV) or *M. smegmatis* expressing PPE2 (Msmeg-PPE2) at 1:10 MOI (multiplicity of infection) for 24 hours. Following infection, the cells were washed with gentamicin to a final concentration of 50 µg/ml to remove extracellular bacteria and these infected cells were induced to differentiate into matured adipocytes. At day 10 post-differentiation, the cells were fixed with 10% paraformaldehyde and stained with Oil Red O. For quantification of Oil Red O stain, the dye was extracted using isopropanol and the absorbance was measured at 490 nm using a spectrophotometer. One-way ANOVA followed by Tukey’s HSD test was carried out. p value of < 0.05 was considered to be significant.

**Figure 2—figure supplement 1.**
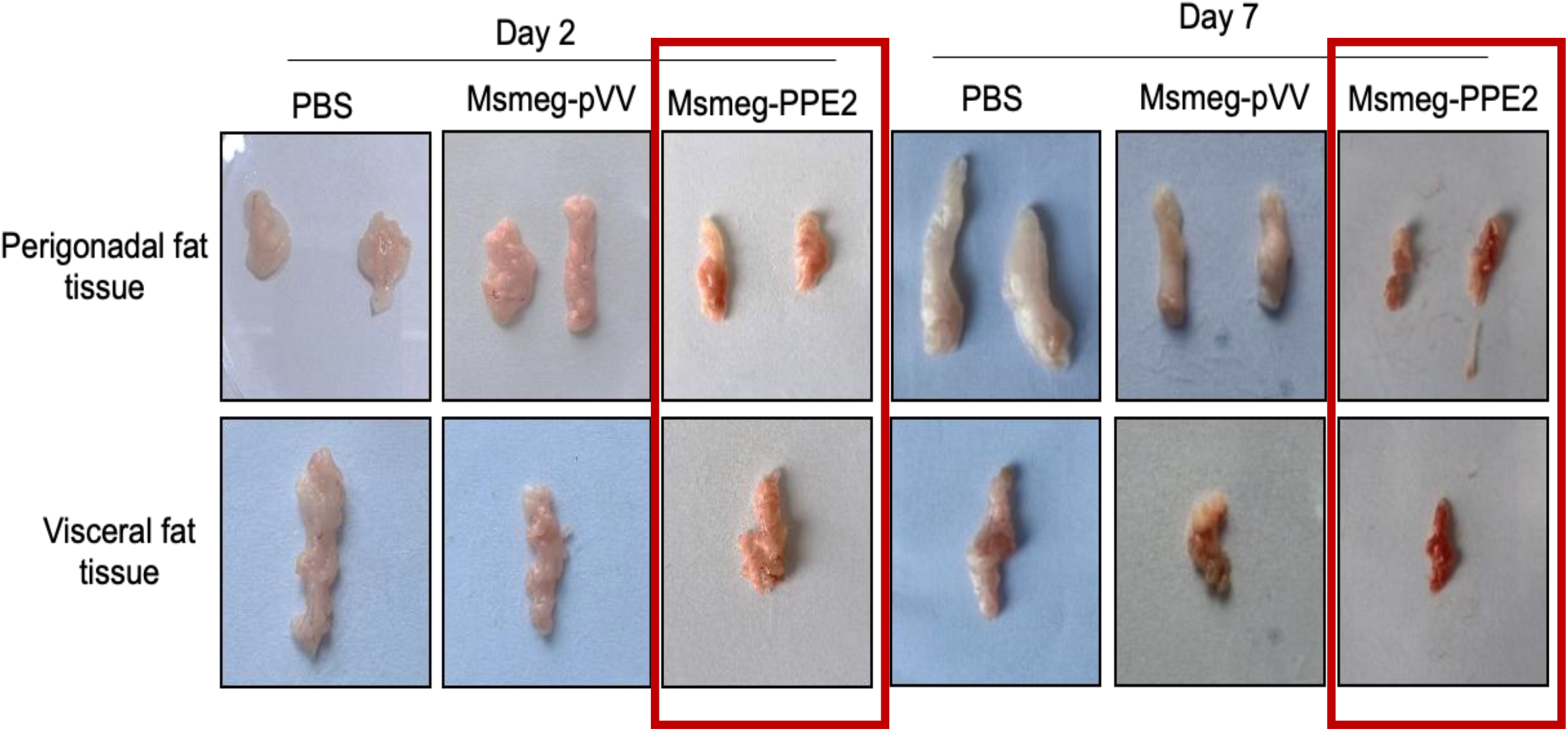
Photographs of perigonadal and visceral fat tissues of mice infected with either Msmeg-pVV or Msmeg-PPE2. Balb/c mice were infected with 100 million of either Msmeg-pVV or Msmeg-PPE2 through intravenous route and were sacrificed on 2^nd^ and 7^th^ day post-infection and the perigonadal and visceral fat tissue were harvested and photographs of the perigonadal and visceral fat tissues from infected mice are shown.

**Figure 2—figure supplement 2.**
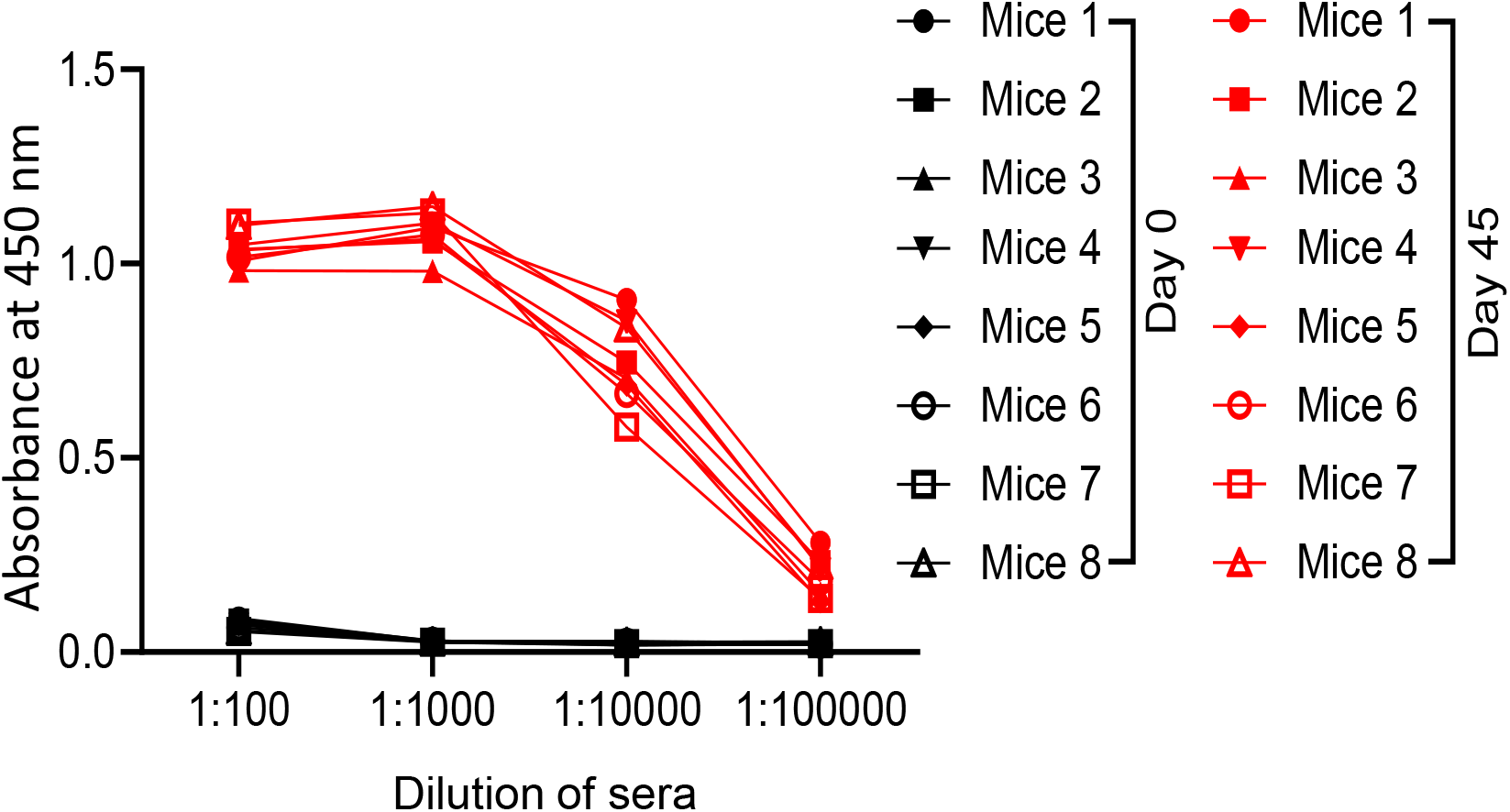
Levels of anti-PPE2 Ab in mice immunized with rPPE2. Balb/c mice (n = 8) were immunized with 100 µg of rPPE2 using incomplete Freund’s adjuvant. Two booster doses were given at 15 days intervals and sera were collected on the 45^th^ day of immunization. Pre-bleed sera were collected at day 0. Sera were diluted and the levels of anti-PPE2 Ab were checked by ELISA. For ELISA, a 96-well microtiter plate was coated with rPPE2 at 1 µg/well diluted in carbonate buffer and incubated overnight at 4°C. After blocking the wells with 2% BSA in PBS, mice sera at various dilutions were added and plate was incubated at 37°C for 1 hour. After washing with PBS-T (0.05% Tween 20 in PBS), the plate was incubated with anti-mouse Ab bound to horseradish peroxidase (HRP) (1:1fig1s1) followed by washing with PBS-T. The plate was then washed and developed with TMB (3, 3’, 5, 5’-Tetramethylbenzidine) substrate (BD Biosciences, USA). The reaction was stopped with 1N H_2_SO_4_ and the absorbance was measured at 450/550 nm in an ELISA microplate reader.

**Figure 2—figure supplement 3.**
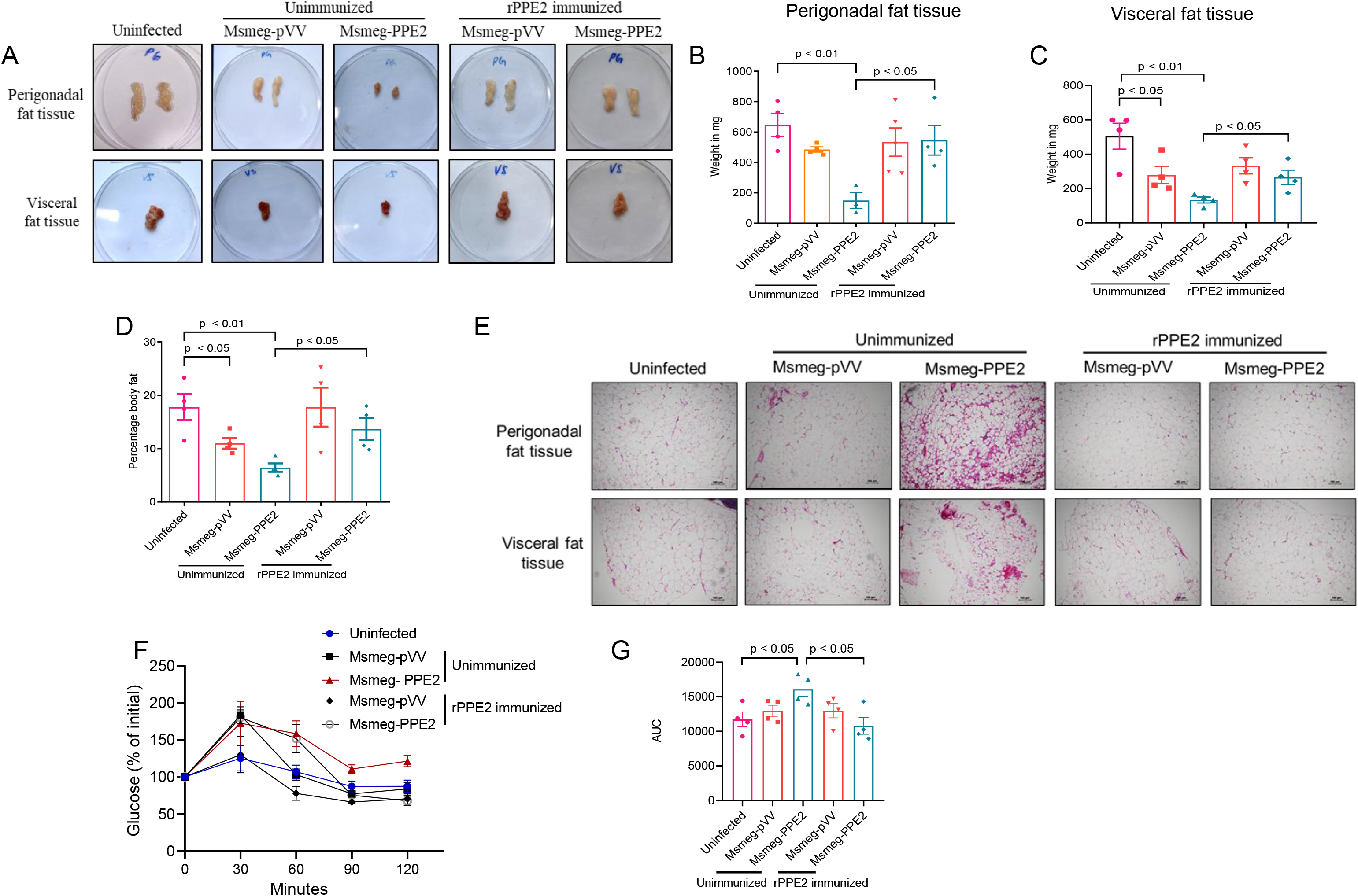
Pre-immunization with rPPE2 protects the mice from detrimental effects of PPE2 on adipose tissue. Mice were immunized with 3 doses of recombinant PPE2 protein with Freund’s incomplete adjuvant at 15 days interval. Next, non-immunized and rPPE2-immunized mice were infected with Msmeg-pVV or Msmeg-PPE2 and a change in fat content was observed after day 7. (**A**) Photographs of perigonadal and visceral fat tissues are shown. Weight of perigonadal (**B**) and visceral (**C**) fat tissues of various groups is recorded. (**D**) Percentage change in total body fat as determined by Dual-energy X-ray Absorptiometry (DEXA) is shown. (**E**) At day 7, sections of perigonadal and visceral fat tissues were prepared and stained with hematoxylin and eosin and photographs of the representative sections were visualized at 40X magnification (scale bar = 100 µm). (**F**) A glucose tolerance test was performed on 7^th^ dpi, and (**G**) the area under the curve (AUC) of glucose concentration is shown. The results shown are mean ± SEM of 4 mice. For Figures B, C, D and G, unpaired Student’s *t* test was carried out. p value of < 0.05 was considered to be significant.

**Figure 4—figure supplement 1.**
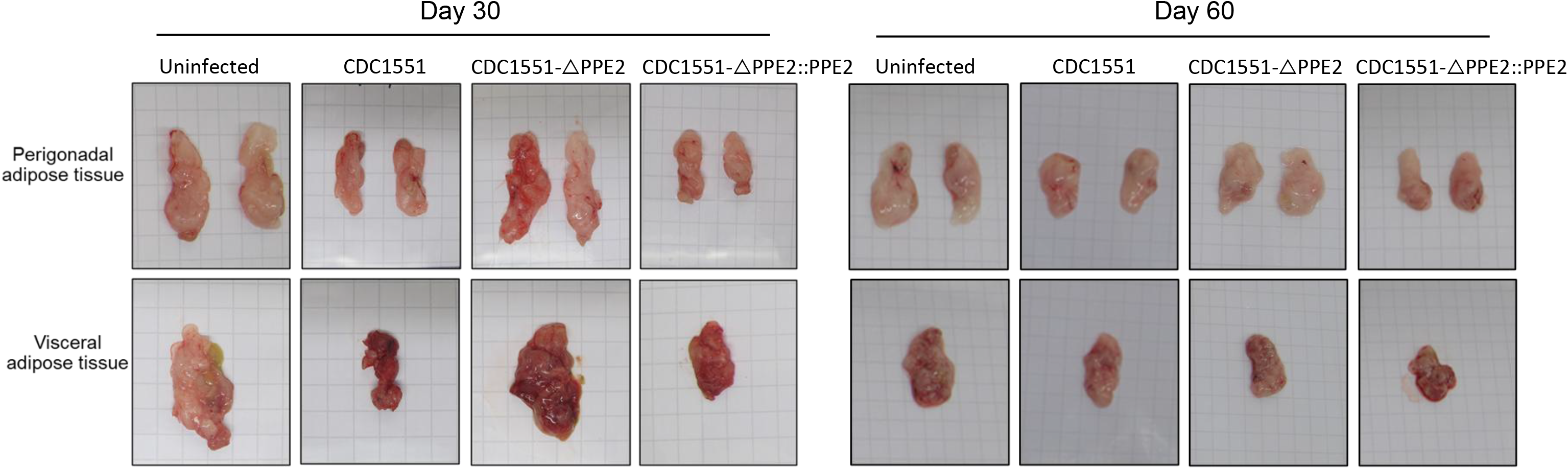
Photographs of perigonadal and visceral fat tissues of mice infected with various strains of *M. tuberculosis*. Balb/c mice were infected *via* aerosol route with wild-type CDC1551 or CDC1551-ΔPPE2 or CDC1551-ΔPPE2::PPE2 and were sacrificed at day 30 or day 60 post-infection. Perigonadal and visceral fat tissues were harvested and photographs of perigonadal and visceral fat tissues are shown.

**Figure 4—figure supplement 2.**
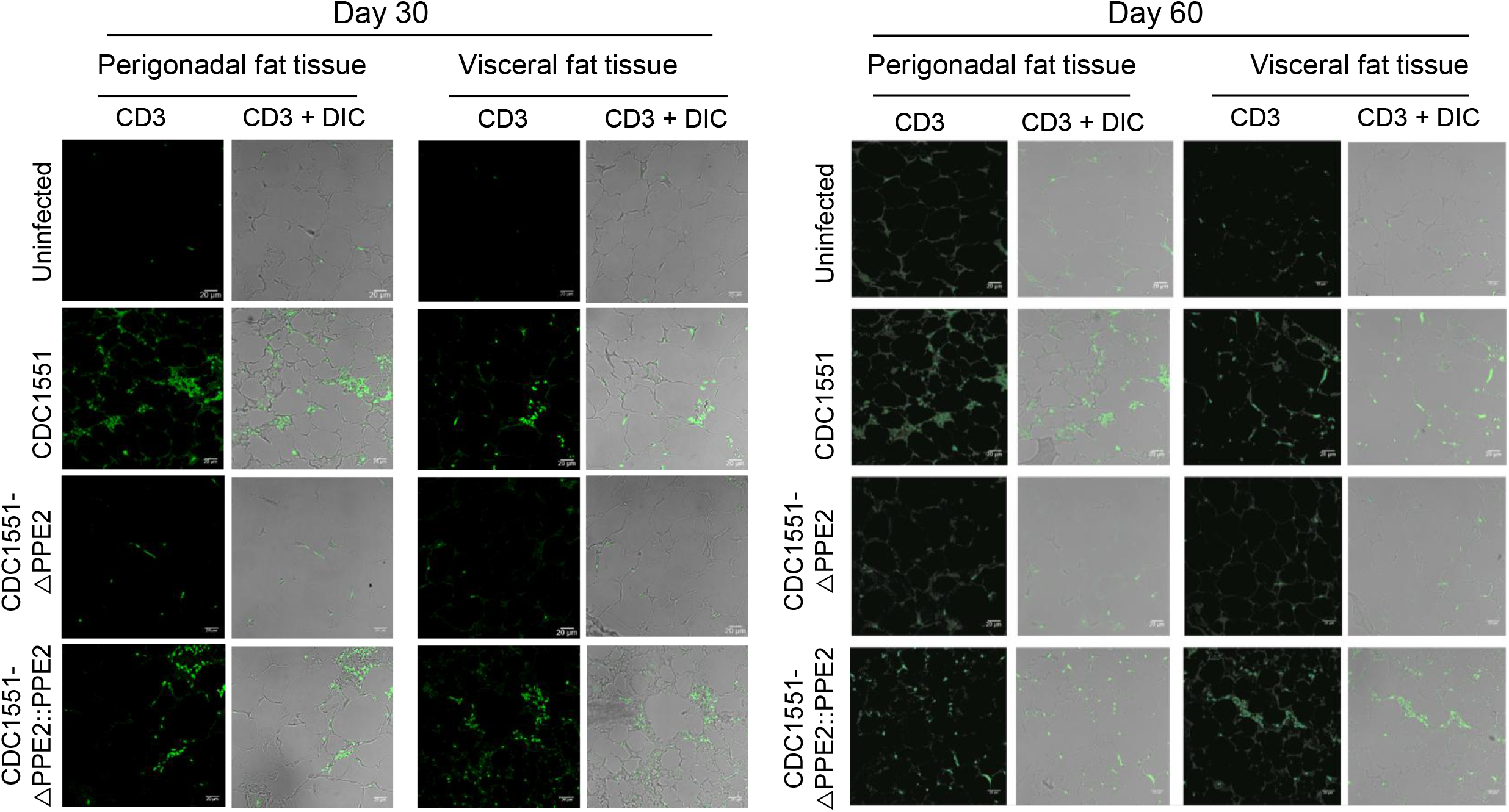
Infiltration of CD3^+^ T-cells in perigonadal and visceral fat tissues. Balb/c mice were infected *via* aerosol route with wild-type *M. tuberculosis* strain CDC1551 or CDC1551-ΔPPE2 or CDC1551-ΔPPE2::PPE2 and were sacrificed at day 30 or day 60 post-infection. Perigonadal and visceral fat tissues were harvested. The tissues were fixed in 4% paraformaldehyde and embedded in paraffin wax and then sectioned using a microtome. The tissue sections were then deparaffinized with 2 changes of xylene followed by sequential hydration with two changes of 100% ethanol for 3 minutes each followed by 95%, 75% and 50% ethanol for 1 minute each. Antigens were retrieved by heating the sections in sodium citrate buffer (10 mM, pH 6) at 95-100°C for 30 minutes, followed by cooling at room temperature. The sections were then premetallized by washing twice with PBS for 2 min each, followed by incubation with 0.1% Triton-X 100 in PBS. Blocking was carried out in PBS containing 5% bovine serum albumin (BSA) for 30 minutes at room temperature. After 3 washes with PBS, the sections were incubated with Pacific Blue™-conjugated anti-mouse CD3 antibody (BioLegend, USA) overnight at 4°C, followed by washing thrice with PBS for 3 minutes each and imaged using a confocal microscope.

**Figure 5—figure supplement 1.**
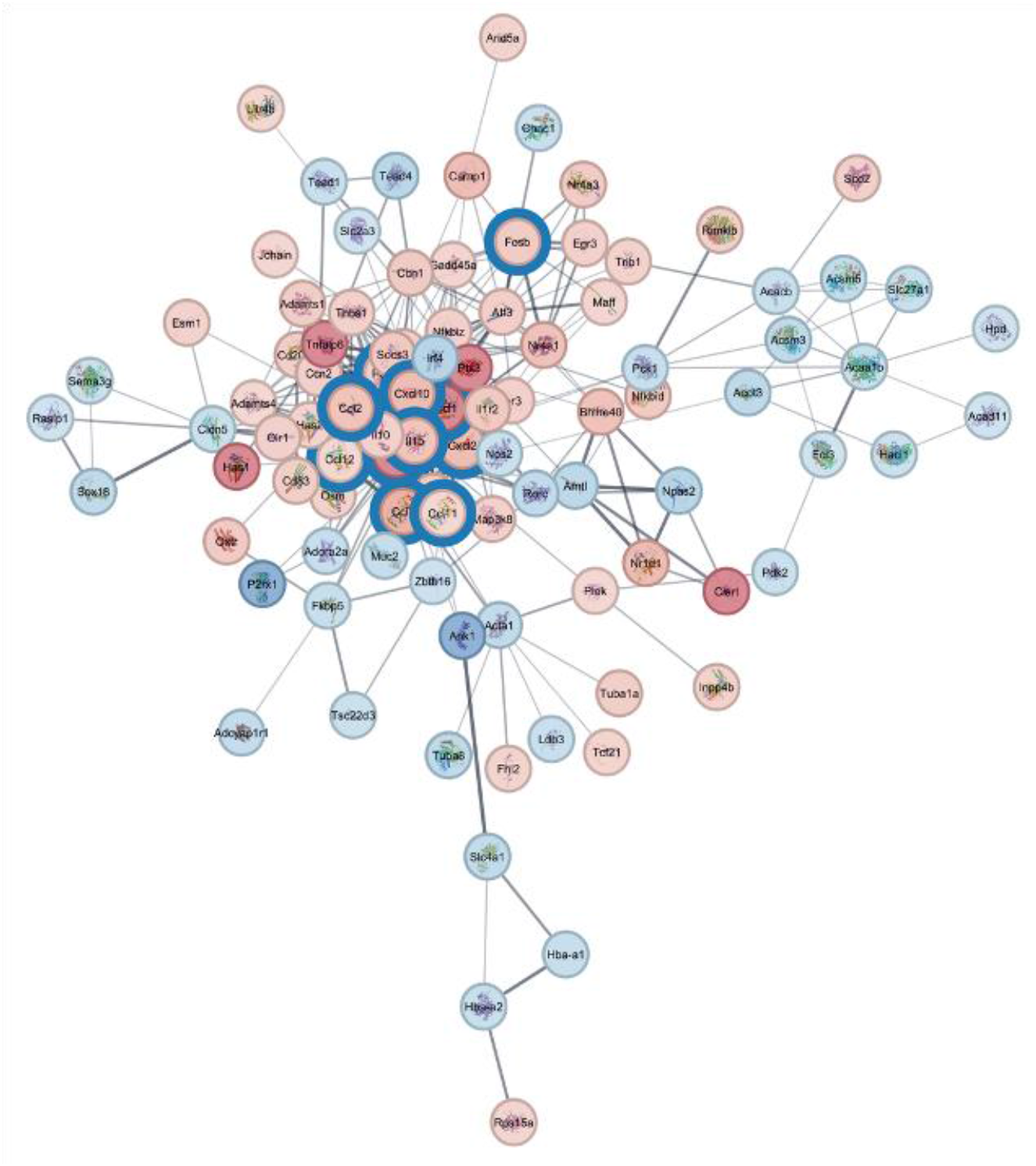
Cystoscape map showing upregulated chemokine map in perigonadal adipose tissue of mice infected with wild-type CDC1551 when compared with adipose tissue of mice infected with CDC1551-ΔPPE2.

**Figure 6—figure supplement 1.**
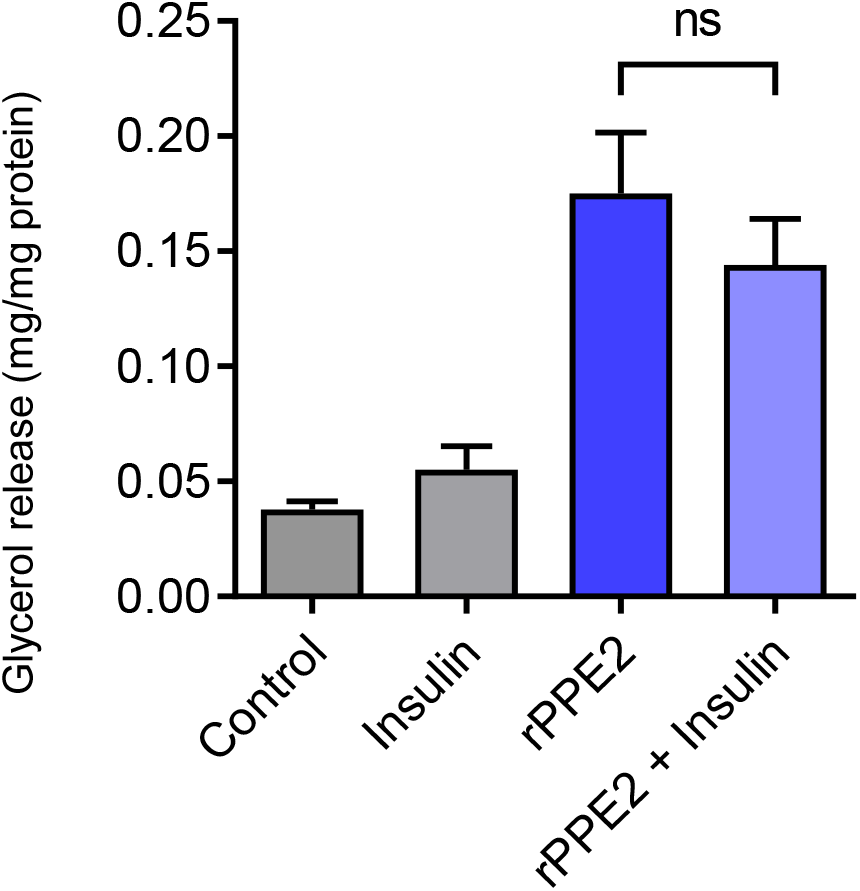
Insulin does not inhibit rPPE2-induced lipolysis. The mature 3T3-L1 adipocytes were treated with rPPE2 protein (3 µg/ml) and after 30 minutes, insulin (1 µg/ml) was added to the medium and lipolysis was measured after 24 hours by estimating free glycerol released in the medium using Free Glycerol Reagent Kit (F6428-40mL, Sigma-Aldrich, USA). Ns = non-significant. Unpaired Student’s *t* test was carried out. p value of < 0.05 was considered to be significant.

**Supplement file 1: Body mass index (BMI) values of patients used in measuring NEFA**

## Notes

### Summary of Updates

1. Statistical analysis details have been added for each figure 2. Beta-actin transcript level was used as an internal control. 3. Figure 5 has now been shifted to Figure 2-Figure Supplement 3. 4. In Figure 3, additional data have been added as Figure 3 A-E. 5. Additional F4/80 macrophage staining data have been added to Figures 3H and 4H-I. 6. Adipocyte hypertrophy has been discussed in the Discussion section. 7. The Discussion section has been shortened. 8. The Key Resources Table has been added to Materials and Methods section. 9. Supplementary File 1, including the body mass index value of TB patients and healthy controls, has been addedton the manuscript. 10. Limitations of the study have been discussed in the Discussion section.

