## Supplementary material for "The PPE2 protein of *Mycobacterium tuberculosis* is responsible for the development of hyperglycemia and insulin resistance during tuberculosis": Figure supplements

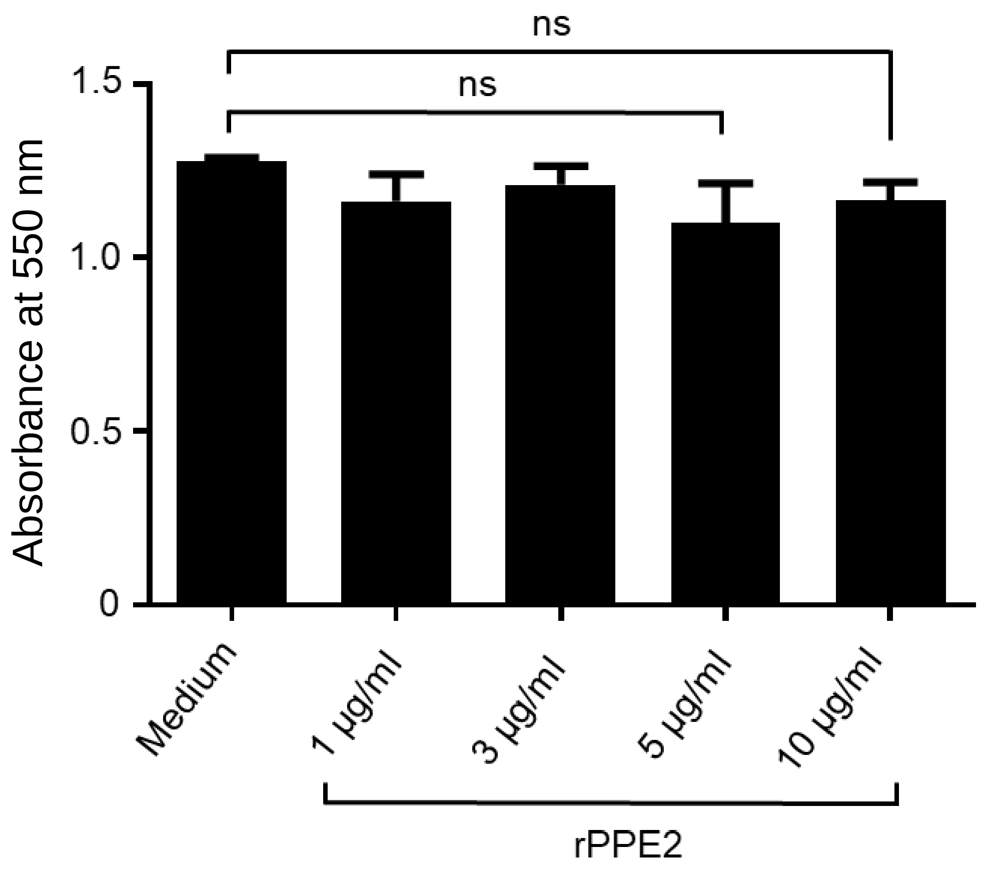


**Figure 1-figure supplement 1. rPPE2 does not cause 3T3-L1 cell toxicity**. 3T3-L1 cells (2 × $10^5$/ml) were treated with different concentrations of rPPE2 for 48 hours and cell cytotoxicity was measured by MTT assay. In brief, MTT (3-(4,5-Dimethylthiazol-2-yl)-2,5-diphenyltetrazolium bromide, Sigma-Aldrich, USA) was added at 1 mg/ml and incubated for 4 hours. The cells were lysed overnight using 100 µl of lysis buffer (20% SDS and 50% dimethyl formamide) and the absorbance was measured at 550 nm. One-way ANOVA followed by Tukey's HSD test was carried out. ns = non-significant, p-value of < 0.05 was considered to be significant.

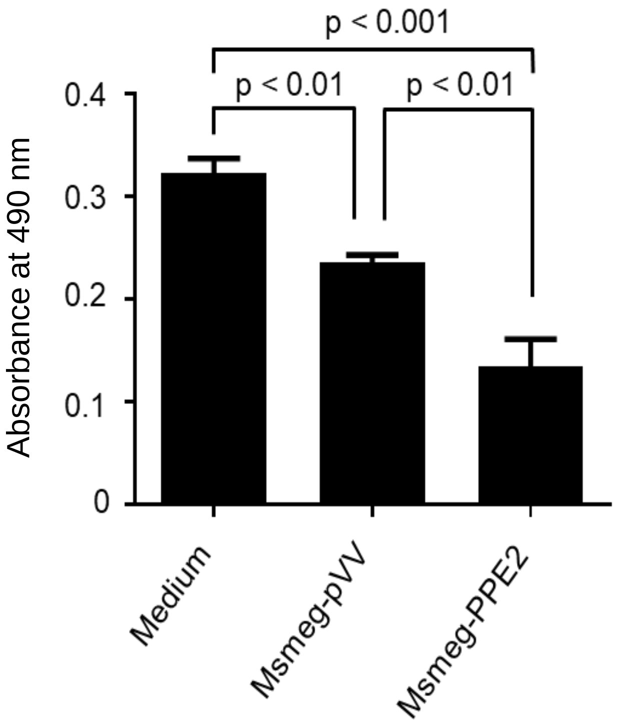


**Figure 1—figure supplement 2. PPE2 inhibits 3T3-L1 adipocyte differentiation during infection.** Undifferentiated 3T3-L1 pre-adipocytes were infected with *M. smegmatis* harboring pVV16 empty vector (Msmeg-pVV) or *M. smegmatis* expressing PPE2 (Msmeg-PPE2) at 1:10 MOI (multiplicity of infection) for 24 hours. Following infection, the cells were washed with gentamicin to a final concentration of 50 µg/ml to remove extracellular bacteria and these infected cells were induced to differentiate into matured adipocytes. At day 10 post-differentiation, the cells were fixed with 10% paraformaldehyde and stained with Oil Red O. For quantification of Oil Red O stain, the dye was extracted using isopropanol and the absorbance was measured at 490 nm using a spectrophotometer. One-way ANOVA followed by Tukey's HSD test was carried out. p value of $< 0.05$ was considered to be significant.

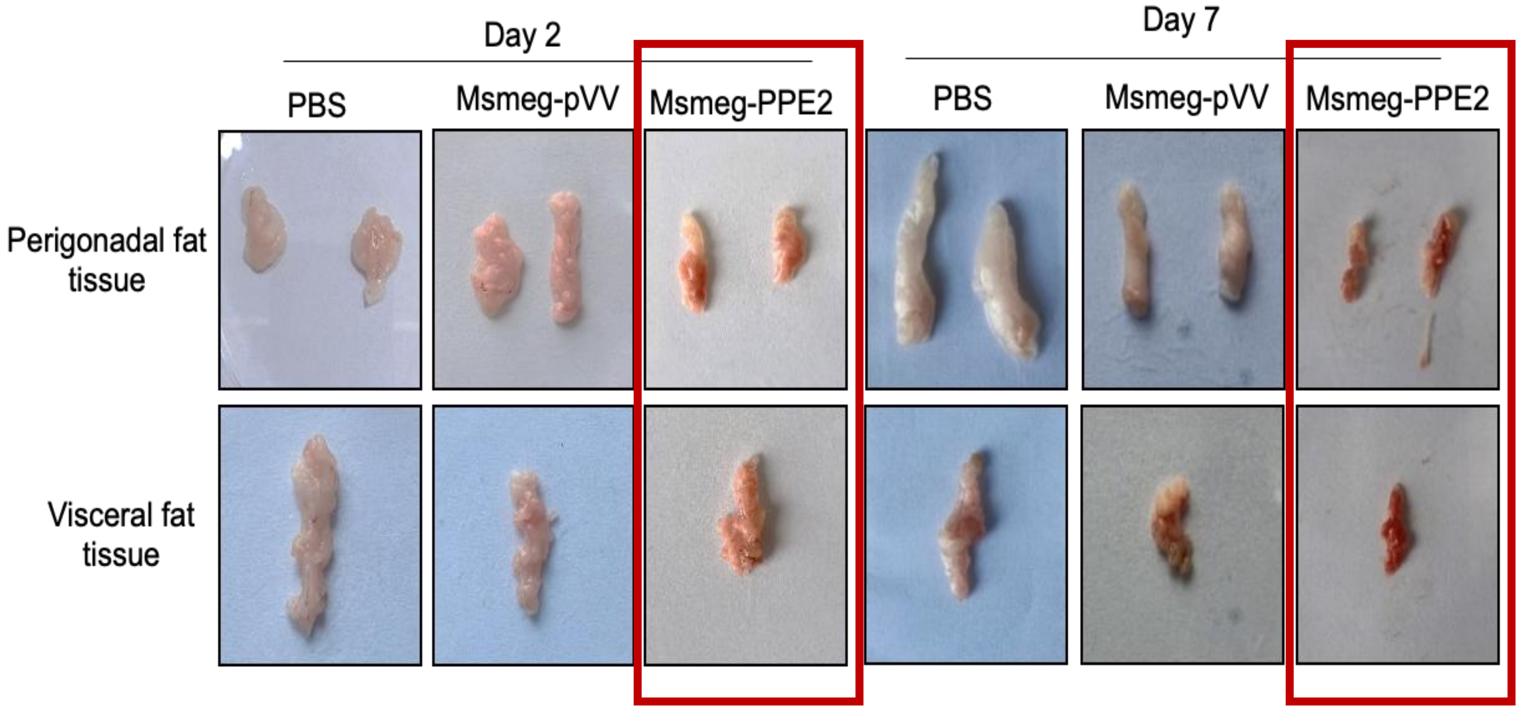


**Figure 2—figure supplement 1. Photographs of perigonadal and visceral fat tissues of mice infected with either Msmeg-pVV or Msmeg-PPE2.** Balb/c mice were infected with 100 million of either Msmeg-pVV or Msmeg-PPE2 through intravenous route and were sacrificed on 2$^{nd}$ and 7$^{th}$ day post-infection and the perigonadal and visceral fat tissue were harvested and photographs of the perigonadal and visceral fat tissues from infected mice are shown.

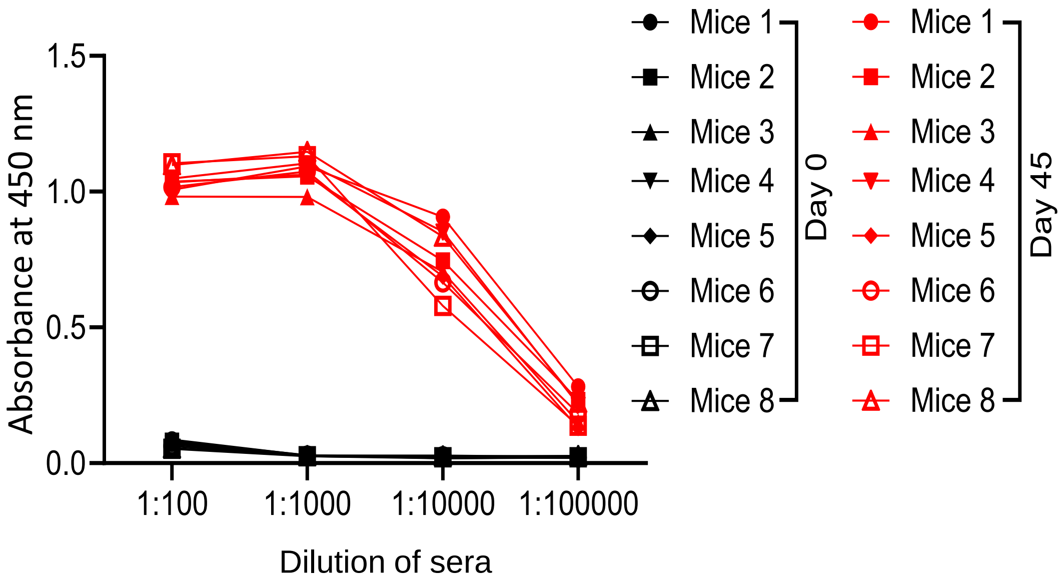


**Figure 2—figure supplement 2. Levels of anti-PPE2 antibody (Ab) in mice immunized with rPPE2.** Balb/c mice (n = 8) were immunized with 100 µg of rPPE2 using incomplete Freund's adjuvant. Two booster doses were given at 15 days intervals and sera were collected on the 45$^{th}$ day of immunization. Pre-bleed sera were collected at day 0. Sera were diluted and the levels of anti-PPE2 Ab were checked by ELISA. For ELISA, a 96-well microtiter plate was coated with rPPE2 protein at 1 µg/well diluted in carbonate buffer and incubated overnight at 4$^{o}$C. After blocking the wells with 2% BSA in PBS, mice sera at various dilutions were added and plate was incubated at 37$^{o}$C for 1 hour. After washing with PBS-T (0.05% Tween 20 in PBS), the plate was incubated with anti-mouse Ab bound to horseradish peroxidase (HRP) (1:10000) followed by washing with PBS-T. The plate was then washed and developed with TMB (3, 3', 5, 5'-Tetramethylbenzidine) substrate (BD Biosciences, USA). The reaction was stopped with 1N $H_2SO_4$ and the absorbance was measured at 450/550 nm in an ELISA microplate reader.

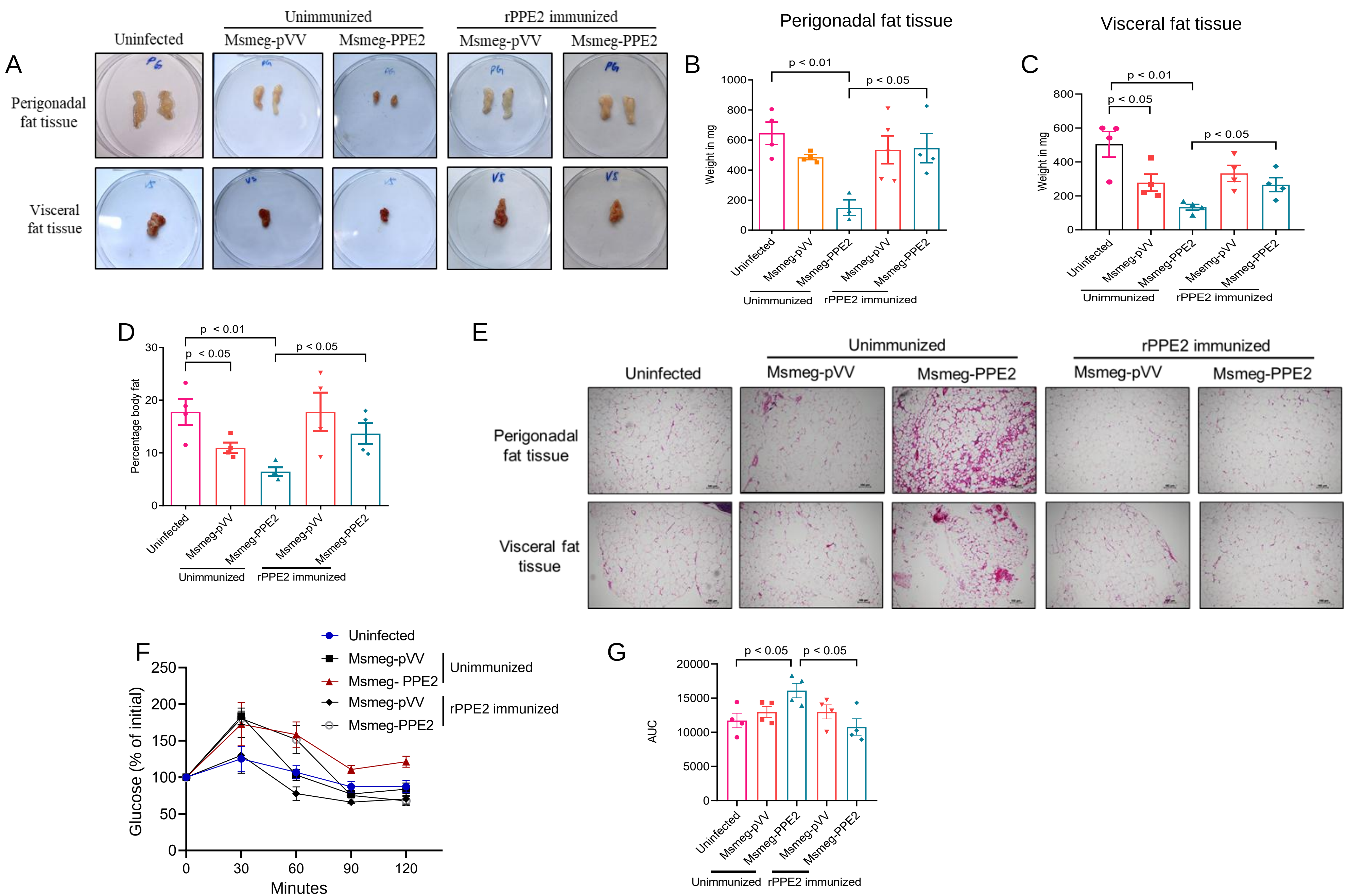

A
Uninfected
Unimmunized
Msmeg-pVV
Msmeg-PPE2
rPPE2 immunized
Msmeg-pVV
Msmeg-PPE2
Perigonadal fat tissue
Visceral fat tissue
B
Perigonadal fat tissue
Weight in mg
p < 0.01
p < 0.05
Uninfected
Msmeg-pVV
Msmeg-PPE2
Msmeg-pVV
Msmeg-PPE2
Unimmunized
rPPE2 immunized
C
Visceral fat tissue
Weight in mg
p < 0.01
p < 0.05
p < 0.05
Uninfected
Msmeg-pVV
Msmeg-PPE2
Msemg-pVV
Msmeg-PPE2
Unimmunized
rPPE2 immunized
D
Percentage body fat
p < 0.01
p < 0.05
p < 0.05
Uninfected
Msmeg-pVV
Msmeg-PPE2
Msmeg-pVV
Msmeg-PPE2
Unimmunized
rPPE2 immunized
E
Uninfected
Unimmunized
Msmeg-pVV
Msmeg-PPE2
rPPE2 immunized
Msmeg-pVV
Msmeg-PPE2
Perigonadal fat tissue
Visceral fat tissue
F
Uninfected
Msmeg-pVV
Msmeg- PPE2
Unimmunized
Msmeg-pVV
Msmeg-PPE2
rPPE2 immunized
Glucose (% of initial)
Minutes
G
AUC
p < 0.05
p < 0.05
Uninfected
Msmeg-pVV
Msmeg-PPE2
Msmeg-pVV
Msmeg-PPE2
Unimmunized
rPPE2 immunized

**Figure 2—figure supplement 3**. **Pre-immunization with rPPE2 protects the mice from detrimental effects of PPE2 on adipose tissue**. Mice were immunized with 3 doses of recombinant PPE2 protein with Freund's incomplete adjuvant at 15 days interval. Next, non-immunized and rPPE2-immunized mice were infected with Msmeg-pVV or Msmeg-PPE2 and a change in fat content was observed after day 7. (**A**) Photographs of perigonadal and visceral fat tissues are shown**.** Weight of perigonadal (**B**) and visceral (**C**) fat tissues of various groups is recorded. (**D**) Percentage change in total body fat as determined by Dual-energy X-ray Absorptiometry (DEXA) is shown. (**E**) At day 7, sections of perigonadal and visceral fat tissues were prepared and stained with hematoxylin and eosin and photographs of the representative sections were visualized at 40X magnification (scale bar = 100 µm). (**F**) A glucose tolerance test was performed on $7^{th}$ dpi and (**G**) Area under the curve (AUC) of glucose concentration is shown. The results shown are mean ± SEM of 4 mice. For Figures B, C, D and G, unpaired Student's *t* test was carried out. p value of $< 0.05$ was considered to be significant

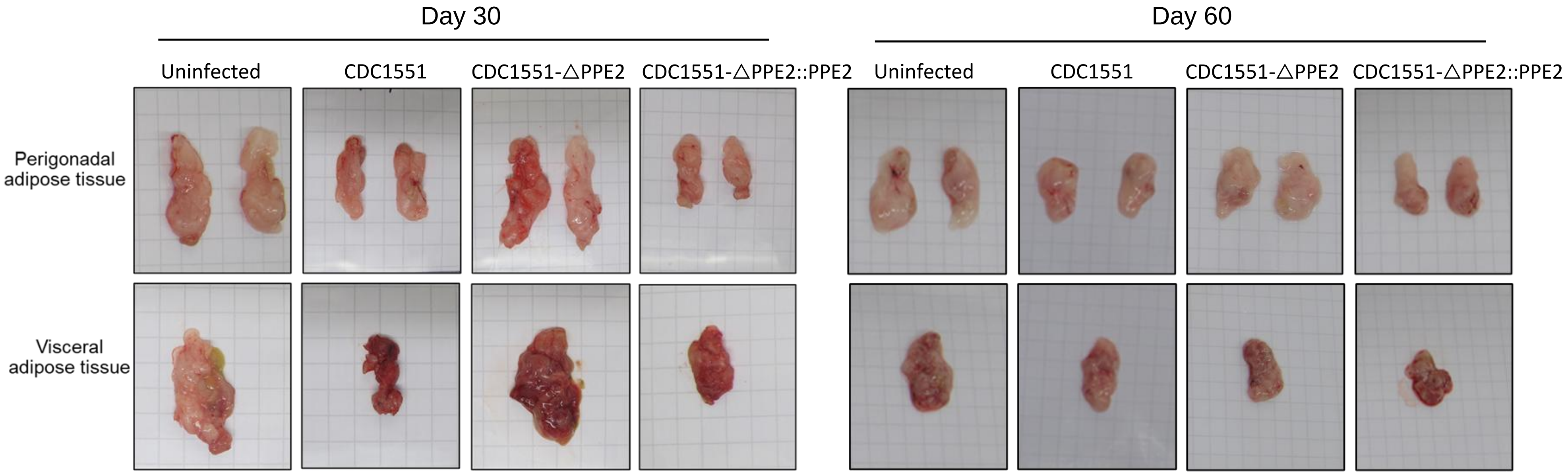


**Figure 4—figure supplement 1. Photographs of perigonadal and visceral fat tissues of mice infected with various strains of *M. tuberculosis*.** Balb/c mice were infected *via* aerosol route with wild-type CDC1551 or CDC1551-ΔPPE2 or CDC1551-ΔPPE2::PPE2 and were sacrificed at day 30 or day 60 post-infection. Perigonadal and visceral fat tissues were harvested and photographs of perigonadal and visceral fat tissues are shown.

.

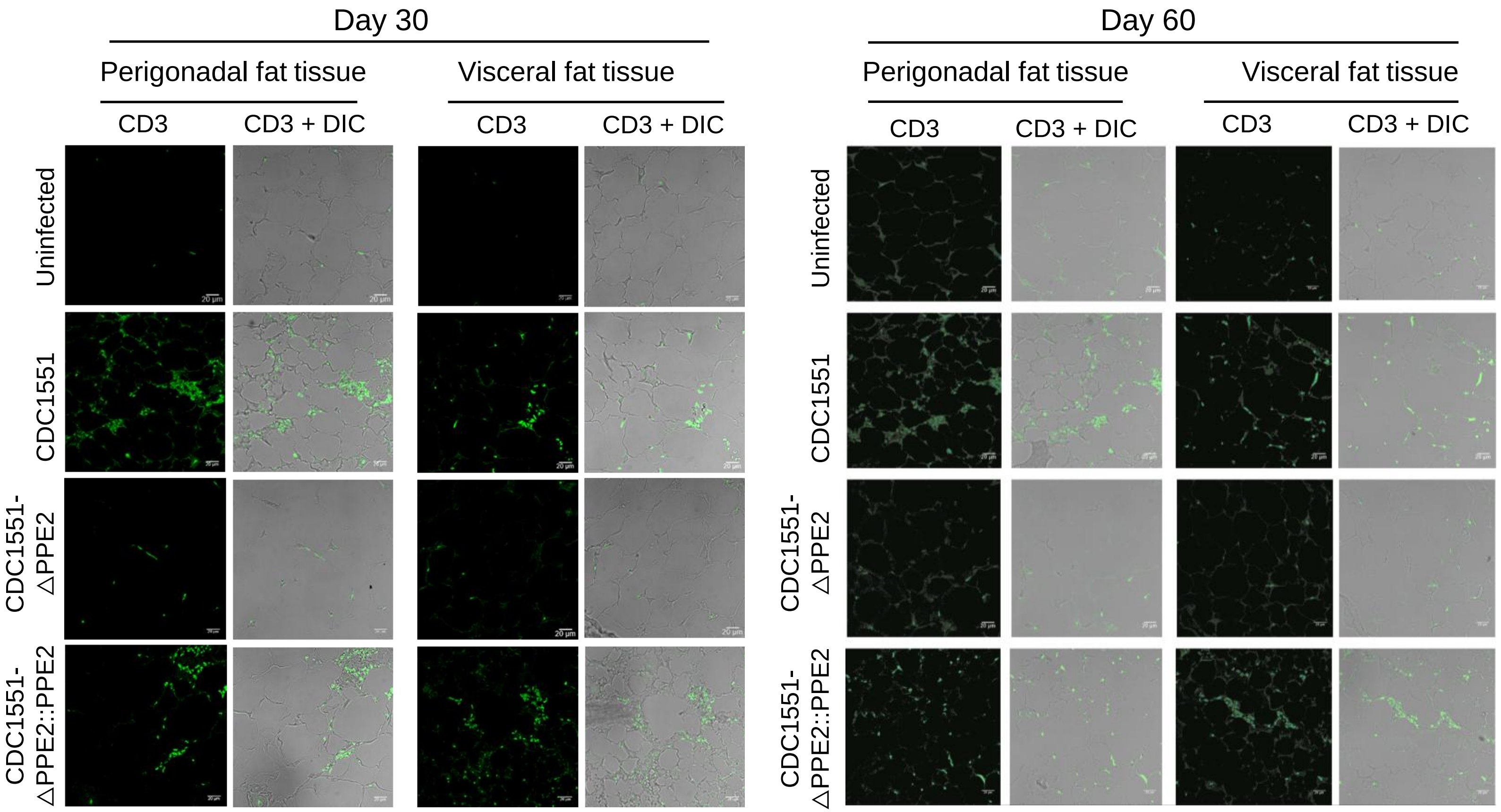


**Figure 4—figure supplement 2. Infiltration of $CD3^+$ T-cells in perigonadal and visceral fat tissues.** Balb/c mice were infected *via* aerosol route with wild-type *M. tuberculosis* strain CDC1551 or CDC1551-ΔPPE2 or CDC1551-ΔPPE2::PPE2 and were sacrificed at day 30 or day 60 post-infection. Perigonadal and visceral fat tissues were harvested. The tissues were fixed in 4% paraformaldehyde and embedded in paraffin wax and then sectioned using a microtome. The tissue sections were then deparaffinized with 2 changes of xylene followed by sequential hydration with two changes of 100% ethanol for 3 minutes each followed by 95%, 75% and 50% ethanol for 1 minute each. Antigens were retrieved by heating the sections in sodium citrate buffer (10 mM, pH 6) at 95-100°C for 30 minutes, followed by cooling at room temperature. The sections were then premetallized by washing twice with PBS for 2 min each, followed by incubation with 0.1% Triton-X 100 in PBS. Blocking was carried out in PBS containing 5% bovine serum albumin (BSA) for 30 minutes at room temperature. After 3 washes with PBS, the sections were incubated with Pacific Blue™-conjugated anti-mouse CD3 antibody (BioLegend, USA) overnight at 4°C, followed by washing thrice with PBS for 3 minutes each and imaged using a confocal microscope.

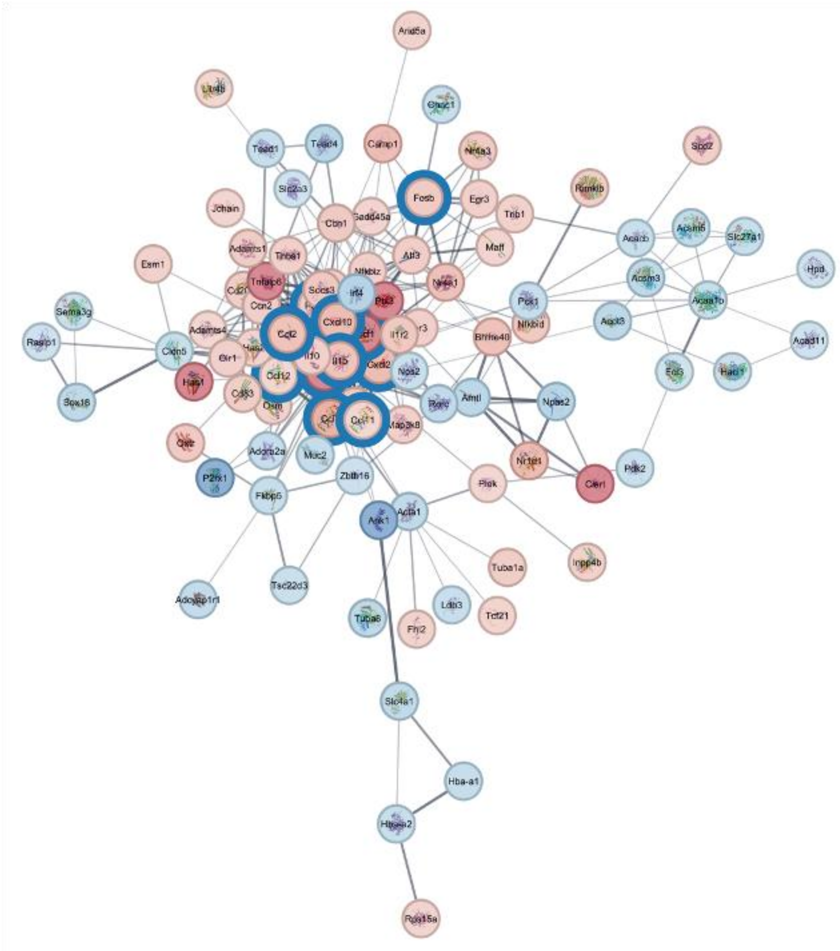

**Figure 5—figure supplement 1.** Cystoscape map showing upregulated chemokine map in perigonadal adipose tissue of mice infected with wild-type CDC1551 when compared with adipose tissue of mice infected with CDC1551-ΔPPE2.

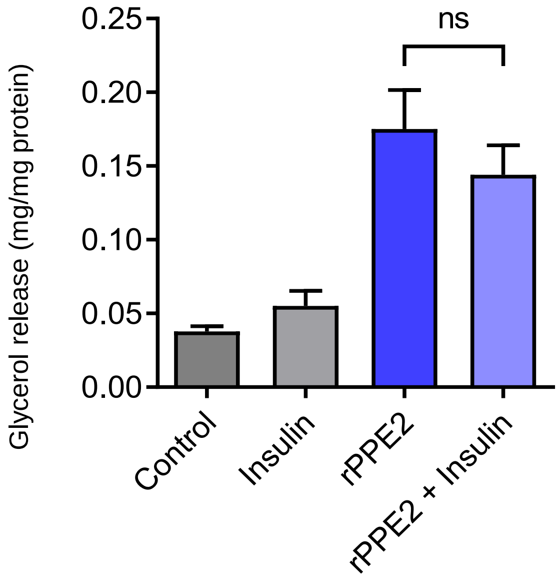


**Figure 6—figure supplement 1. Insulin does not inhibit rPPE2-induced lipolysis.** The mature 3T3-L1 adipocytes were treated with rPPE2 protein (3 µg/ml) and after 30 minutes, insulin (1 µg/ml) was added to the medium and lipolysis was measured after 24 hours by estimating free glycerol released in the medium using Free Glycerol Reagent Kit (F6428-40mL, Sigma-Aldrich, USA). ns = non-significant. Unpaired Student's *t* test was carried out. p value of $< 0.05$ was considered to be significant.
