## Supplement file 1 for "The PPE2 protein of *Mycobacterium tuberculosis* is responsible for the development of hyperglycemia and insulin resistance during tuberculosis"

| **S. No** | **Age (years)** | **Date of sample collection** | **Sex (M/F)** | **Weight (kg)** | **Height (cm)** | **BMI (kg m-2)** | **Alcohol Use (Yes/No)** | **Type** |
| --- | --- | --- | --- | --- | --- | --- | --- | --- |
| 1 | 25 | 28-03-2023 | MALE | 54 | 168 | 19.13 | yes | **Pulmonary TB** |
| 2 | 22 | 27-04-2023 | MALE | 44 | 178 | 13.89 | yes |  |
| 3 | 18 | 12-05-2023 | FEMALE | 25 | 165 | 9.18 | no |  |
| 4 | 65 | 12-05-2023 | FEMALE | 25 | 168 | 8.86 | yes-ex |  |
| 5 | 75 | 18-05-2023 | MALE | 59 | 162 | 22.48 | yes |  |
| 6 | 38 | 18-05-2023 | MALE | 42 | 158 | 16.82 | yes |  |
| 7 | 27 | 18-05-2023 | MALE | 39 | 172 | 13.18 | yes |  |
| 8 | 30 | 22-05-2023 | MALE | 40 | 162 | 15.24 | no |  |
| 9 | 60 | 22-05-2023 | MALE | 38 | 153 | 16.23 | yes |  |
| 10 | 23 | 07-07-2023 | FEMALE | 32 | 154 | 13.49 | no |  |
| 11 | 28 | 07-07-2023 | MALE | 59 | 170 | 20.42 | yes |  |
| 12 | 60 | 07-07-2023 | MALE | 68 | 170 | 23.53 | yes |  |
| 13 | 65 | 27-07-2023 | MALE | 48 | 162 | 18.29 | no |  |
| 14 | 17 | 16-08-2023 | FEMALE | 45 | 160 | 17.58 | no |  |
| 15 | 50 | 16-08-2023 | MALE | 42 | 172 | 14.20 | yes |  |
| 16 | 50 | 13-09-2023 | MALE | 45 | 158 | 18.03 | yes |  |
| 17 | 57 | 04-10-2023 | MALE | 50 | 168 | 17.72 | yes |  |
| 18 | 58 | 04-10-2023 | MALE | 60 | 175 | 19.59 | yes |  |
| 19 | 44 | 04-10-2023 | MALE | 55 | 168 | 19.49 | yes |  |
| 20 | 45 | 05-10-2023 | MALE | 45 | 156 | 18.49 | yes |  |
| 21 | 58 | 05-10-2023 | MALE | 38 | 149 | 17.12 | yes |  |
| 22 | 57 | 25-10-2023 | MALE | 58 | 155 | 24.14 | yes |  |
| 23 | 53 | 25-10-2023 | MALE | 45 | 173 | 15.04 | yes |  |
| 24 | 45 | 28-10-2023 | FEMALE | 55 | 149 | 24.77 | no |  |
| 25 | 33 | 28-10-2023 | MALE | 50 | 177 | 15.96 | no |  |
| 26 | 22 | 06-11-2023 | FEMALE | 42 | 168 | 14.88 | no |  |
| 27 | 38 | 06-11-2023 | MALE | 56 | 158 | 22.43 | yes |  |
| 28 | 45 | 06-11-2023 | MALE | 42 | 178 | 13.26 | yes |  |
| 29 | 60 | 06-11-2023 | MALE | 39 | 170 | 13.49 | no |  |
| 30 | 24 | 14-11-2023 | Female | 40 | 150 | 17.78 | no |  |
| 31 | 48 | 14-11-2023 | Male | 42 | 176 | 13.56 | yes |  |
| 32 | 51 | 14-11-2023 | Male | 58 | 172 | 19.61 | yes |  |
| 33 | 40 | 14-11-2023 | Male | 48 | 178 | 15.15 | yes |  |
| 34 | 25 | 14-11-2023 | Female | 50 | 162 | 19.05 | no |  |
| 35 | 38 | 15-11-2023 | Male | 55 | 158 | 22.03 | yes |  |
| 36 | 44 | 16-04-2018 | Male | 40 | 140 | 20.41 | Unknown |  |
| 37 | 43 | 31-08-2019 | Male | 53 | 152 | 22.94 | Unknown |  |
| 38 | 23 | 16-05-2020 | Male | 40 | 168 | 14.17 | Unknown |  |
| 39 | 21 | 01-05-2018 | Male | 35 | 142 | 17.36 | Unknown |  |
| 40 | 30 | 12-04-2019 | Male | 38 | 148 | 17.35 | Unknown |  |
| 41 | 19 | 23-06-2019 | Female | 35 | 155 | 14.57 | Unknown |  |
| 42 | 20 | 24-09-2021 | Female | 38 | 153 | 16.23 | Unknown |  |
| 43 | 45 | 30-10-2021 | Male | 65 | 160 | 25.00 | Unknown |  |
| 44 | 47 | 25-11-2021 | Male | 76 | 178 | 24.00 | Unknown |  |
| 45 | 21 | 10-12-2021 | Female | 33 | 146 | 15.50 | Unknown |  |
| 46 | 25 | 27-12-2021 | Female | 40 | 151 | 17.50 | Unknown |  |
| 47 | 40 | 31-05-2022 | Male | 49 | 152 | 21.20 | Unknown |  |
| 48 | 47 | 02-03-2023 | MALE | 48 | 162 | 18.29 | no |  |
| 49 | 39 | 06-03-2023 | MALE | 35 | 160 | 13.67 | yes |  |
| 50 | 51 | 05-10-2023 | Male | 60.2 | 172 | 20.35 | yes |  |
| 51 | 27 | 05-10-2023 | Male | 62 | 170 | 21.45 | no |  |

**Healthy Control**

| 1 | 36 | 15-03-2022 | Female | 60 | 165 | 22.04 | NO | **Healthy control** |
| --- | --- | --- | --- | --- | --- | --- | --- | --- |
| 2 | 35 | 22-04-2022 | Male | 84 | 182 | 25.36 | YES |  |
| 3 | 39 | 26-07-2023 | Female | 55 | 158 | 22.03 | NO |  |
| 4 | 39 | 26-07-2023 | Female | 80 | 176 | 25.83 | NO |  |
| 5 | 21 | 26-07-2023 | Female | 40 | 154 | 16.87 | NO |  |
| 6 | 16 | 26-07-2023 | Male | 63 | 167 | 22.59 | NO |  |
| 7 | 16 | 26-07-2023 | Male | 59 | 180 | 18.21 | NO |  |
| 8 | 26 | 23-08-2023 | Male | 71 | 160 | 27.73 | YES |  |
| 9 | 36 | 10.10.2023 | Male | 78 | 171 | 26.67 | NO |  |
| 10 | 36 | 10.10.2023 | Male | 69 | 168 | 24.45 | Yes |  |
| 11 | 32 | 10.10.2023 | Female | 55 | 160 | 21.48 | NO |  |
| 12 | 30 | 10.10.2023 | Female | 52 | 161 | 20.06 | NO |  |
| 13 | 34 | 10.10.2023 | Male | 79 | 167 | 28.33 | Yes |  |
| 14 | 34 | 10.10.2023 | Male | 78 | 169 | 27.31 | Yes |  |
